# ALG-2 interacting protein-X (Alix) is required for activity-dependent bulk endocytosis at brain synapses

**DOI:** 10.1101/2020.07.20.211854

**Authors:** Marine H. Laporte, Kwang Il Chi, Laura C. Caudal, Na Zhao, Marta Rolland, José Martinez-Hernandez, Magalie Martineau, Christine Chatellard, Eric Denarier, Vincent Mercier, Florent Lemaître, Béatrice Blot, Eve Moutaux, Maxime Cazorla, David Perrais, Fabien Lanté, Sandrine Fraboulet, Fiona J. Hemming, Frank Kirchhoff, Rémy Sadoul

## Abstract

In chemical synapses undergoing high frequency stimulation, vesicle components can be retrieved from the plasma membrane via a clathrin-independent process called activity dependent bulk endocytosis (ADBE). Alix (ALG-2 interacting protein X)/ PDCD6IP) is an adaptor protein binding to ESCRT and endophilin-A proteins which is required for clathrin-independent endocytosis in fibroblasts. Alix is expressed in neurons and concentrates at synapses during epileptic seizures. Here, we used cultured neurons to show that Alix is recruited to presynapses where it interacts with, and concentrates endophilin-A during conditions triggering ADBE. Using Alix knockout (ko) neurons we showed that this recruitment, which requires interaction with the calcium-binding protein ALG-2, is necessary for ABDE. We also found that presynaptic compartments of Alix ko hippocampi display subtle morphological defects compatible with flawed synaptic activity and plasticity detected electrophysiologically. Furthermore, mice lacking Alix in the forebrain undergo less seizures during kainate-induced status epilepticus and reduced propagation of the epileptiform activity. These results thus show that impairment of ADBE due to the lack of neuronal Alix alters synaptic recovery during physiological or pathological repeated stimulations.

## Introduction

Neuronal communication in mammalian brain relies heavily on the activity-dependent release of chemical neurotransmitters from presynaptic boutons. Following fusion of synaptic vesicles (SV) with the presynaptic membrane, SV lipids and proteins are retrieved by endocytosis. Endocytosis avoids detrimental increase in the plasma membrane surface and allows recycling of the SV components to replenish the SV pool (Gan and Watanabe, 2018). At moderate levels of stimulation, retrieval of membrane involves clathrin-mediated (CME) and clathrin-independent endocytosis (CIE), in proportions which are still highly debated (Chanaday and Kavalali, 2018). Moreover, long-lasting high-frequency stimulations also lead to the clathrin-independent internalization of large stretches of presynaptic membranes. This calcium-dependent process, first discovered at the amphibian neuromuscular junction (Miller and Heuser, 1984) is referred to as activity-dependent bulk endocytosis (ADBE). It is meant to avoid abnormal increase of the synaptic bouton surface and to allow replenishment of SVs during sustained synaptic stimulations (Cheung and Cousin, 2012; Marxen et al., 1999). Thus, ADBE has been suggested to play key regulatory roles in physiological or pathological events like epilepsy, which are triggered and sustained by high frequency neuronal activity. However, decrypting the physiological role of ADBE has been hindered by the lack of identified molecules that are both specific and essential to this endocytosis mode.

We have recently demonstrated that ALG-2 interacting protein-X (Alix) is essential for clathrin-independent bulk endocytosis in fibroblasts (Mercier et al., 2016). Alix ko mice have normally organized but smaller brains (Campos et al., 2016; Laporte et al., 2017), a phenotype linked with an alteration of CIE in developing neurons (Laporte et al., 2017). In the adult brain, Alix is ubiquitously expressed but concentrates at hippocampal presynaptic terminals during epileptic seizures (Hemming et al., 2004). Alix is a cytosolic protein first identified through its calcium-dependent binding to the penta-EF-hand protein ALG-2 (apoptosis-linked gene 2) which helps opening and activating the protein (Chatellard-Causse et al., 2002; Missotten et al., 1999; Sadoul, 2006). Its activation leads to interaction with membranes as reported in the case of plasma membrane wounds, where ALG-2 binds to inflowing calcium and helps recruiting Alix to the membrane where the latter organizes repair (Scheffer et al., 2014). Alix also interacts with lipids (Bissig et al., 2013) and with several membrane modifying proteins among which endophilin-A proteins (A1, A2 and A3) (Chatellard-Causse et al., 2002). These cytoplasmic proteins that contain an N-BAR (Bin/Amphiphysin/Rvs) domain capable of sensing and generating membrane curvature (Kjaerulff et al., 2011), are major actors of CME at synapses (Gad et al., 2000; Ringstad et al., 1999). They also drive CIE in fibroblasts (Boucrot et al., 2014; Renard et al., 2014) and were shown to control the fast mode of CIE at ribbon synapses (Llobet et al., 2011) as well as in hippocampal neurons (Kononenko et al., 2014; Watanabe et al., 2018).

The role of Alix in bulk endocytosis in fibroblasts, its capacity to interact with endophilin-A and to be recruited by calcium at membranes, together with its increased concentration at hippocampal synapses during kainate-induced seizures, brought us to test its possible function in ADBE.

Using cultured wildtype (wt) neurons, we now bring evidence that sustained synaptic activity leads to calcium-dependent recruitment of ALG-2 at synapses. ALG-2 in turn interacts with Alix, which binds and concentrates endophilin-A. This protein complex is necessary for ADBE, which is selectively impaired in Alix ko neurons. We also report that synapse morphology and function are both altered in Alix ko brains. This finding correlates with the modifications in kinetics of synaptic recovery following prolonged stimulation that we detected in hippocampal slices of mice in which Alix was selectively deleted in forebrain neurons. In these mice, the number of seizures during status epilepticus induced by intracortical kainate injections was reduced as well as the propagation of epileptiform activity to the contralateral side of injection. Thus, our results show that some molecular mechanisms involved in ADBE may also be involved in certain aspects of synaptic plasticity such as short-term synaptic depression and recurrence of epileptic seizures.

## Material and methods

### Plasmids

Endophilin A2-mCherry was obtained by subcloning (In-Fusion Cloning kit, Clontech) endophilin A2 cDNA into a pmCherry-N1 vector (Clontech). GFP-ALG-2 was obtained by performing a reverse mutagenesis (Quick change II site directed mutagenesis kit, Stratagene) on a GFP-hALG-2 Y180A (a generous gift from Masatoshi Maki) to acquire GFP-hALG-2wt. hALG-2 E47A-E114A cDNA was kindly provided by Masatoshi Maki (Shibata et al., 2004) and was subcloned into a pEGFP-C1 vector (Clontech) to obtain GFP-hALG-2 E47A-E114A (GFP-ALG-2ΔCa).

All constructs containing Alix cDNA (wt or mutants) were obtained by subcloning the relevant cDNAs from pCI vectors harbouring Alix cDNA or its mutants. Alix I212D (AlixΔCHMP4B) and AlixΔPGY (AlixΔALG-2) cDNAs in pCI were generated by mutagenesis (Quick change II site directed mutagenesis kit, Stratagene) and Alix R757E (AlixΔendo) by in-fusion cloning, using the oligos given below.

mCherry-2Xflag-mAlix wt (mCherry-Alix) was obtained by subcloning 2xflag-mAlix wt cDNA into a pmCherry-C1 vector (Clontech). Alix-YFP was obtained by subcloning wild type Alix cDNA into a pEYFP-N1 vector (Clontech). GFP-flag-Alix (GFP-Alix) was described in (Mercier et al., 2016). GFP-Alix and its mutant forms (GFP-Alix R757E, GFP-AlixΔPGY) were obtained by subcloning the various cDNAs into a pEGFP-C1 vector (Clontech). DNA constructs used for the rescue experiments were prepared in two steps. First, IRES2-GFP cDNA was subcloned into pSIN lentiviral vector (kindly provided by F. Saudou) by using pIRES2-GFP (Clontech) as a template. Then the various cDNAs were subcloned into pSIN-IRES2-GFP.

Oligos used to generate mutants:

Alix I212D

sense: 5’-AAGATGAAAG ATGCCGACAT AGCTAAGCTG-3’

antisense: 5’-CAGCTTAGCT ATGTCGGCAT CTTTCATCTT -3’

Alix R757E

sense: 5’-CAGCCGAGCC TCCACCTCCT GTGCTTCCTG -3’

antisense: 5’-GAGGCTCGGC TGGAGGCTGG GGCTTAGCAG-3’

AlixΔPGY

sense: 5’-GCCACAGGCT CAGGGATGCC AAATGCCCAT GC-3’

antisense: 5’-GCATGGGCAT TTGGCATCCC TGAGCCTGTG GC -3’.

### Antibodies

| Antibody | Supplier (reference) | Species (type) | Dilution |
| --- | --- | --- | --- |
| Anti-Actin | Millipore (MAB1501R) | Mouse (monoclonal) | 1/10000 |
| Anti-Alix | Covalab (ab0204) | Rabbit (polyclonal) | 1/10000 |
| Anti GFAP | Abcam | Goat (polyclonal) | 1/1000 |
| Anti Iba1 | Wako | Rabbit (polyclonal) | 1/500 |
| Anti-phospho-p44/42<br>MAPK (T202/Y204) | Cell signaling (9106) | Mouse (monoclonal) | 1/1000 |
| Anti-PSD95 | NeuroMab (73-028) | Mouse (monoclonal) | 1/2000 |
| Anti-PSD95 | Millipore (MAB1598) | Mouse (monoclonal) | 1/500 |
| Anti-Synapsin-1 | Millipore (AB1543P) | Rabbit (polyclonal) | 1/1000 |
| Anti-Synaptophysin | Merck Millipore (MAB5258) | Mouse (monoclonal) | 1/5000 |
| Anti-Flag | Sigma-Aldrich (F3165) | Mouse (monoclonal) | 1/1000 |
| Anti-Flag | Sigma-Aldrich (F7425) | Rabbit (polyclonal) | 1/1000 |
| Anti-Mouse HRP | Jackson ImmunoResearch (115-035-166) | Goat (polyclonal) | 1/5000 |
| Anti-Rabbit HRP | Jackson ImmunoResearch (115-035-044) | Goat (polyclonal) | 1/5000 |
| Anti-Mouse Alexa Fluor<br>488 | Invitrogen (A-11029) | Goat (polyclonal) | 1/1000 |
| Anti-Mouse Alexa Fluor<br>594 | Invitrogen (A-11032) | Goat (polyclonal) | 1/1000 |
| Anti-Mouse Cy5 | Jackson ImmunoResearch (115-175-146) | Goat (polyclonal) | 1/500 |
| Anti-Rabbit Alexa Fluor<br>488 | Invitrogen (A-11034) | Goat (polyclonal) | 1/1000 |
| Anti-Rabbit Alexa Fluor<br>594 | Invitrogen (A-11037) | Goat (polyclonal) | 1/1000 |
| Anti-Rabbit Cy5 | Jackson ImmunoResearch (111-175-144) | Goat (polyclonal) | 1/1000 |

### Animals

Animals were handled and killed in conformity with European law and internal regulations of INSERM. Pregnant Oncins France souche A (OFA; a substrain of Sprague Dawley) rats (Charles River) were used for rat neuronal cultures. Alix ko C57BL/6 mouse pups (Laporte et al., 2017; Mercier et al., 2016) and their control littermates referred to thereafter as Alix wt, were also used for primary neuronal culture. Transgenic mice were held at the animal facility of the Grenoble Institute for Neurosciences and fed *ad libitum*. All animals were held at a twelve-hour light/dark cycle. One-to two-month-old Alix wt and Alix ko mice were used for electrophysiological extracellular recordings, histochemistry and electron microscopy studies.

Mice were anesthetized by intraperitoneal injection of 0.1 ml sodium pentobarbital (5.6% w/v; CEVA Santé Animale) and treated as described in the corresponding sub-headings of the material and methods section.

Emx1^IREScre^/Alix^fl/fl^ and Emx1^IREScre^ control mice, thereafter referred to control and Alix cko, were used for the kainate injection epilepsy model and the electrophysiological. 8-11 weeks old mice were anesthetized under a mixture of 2 % isoflurane, 47.5 % O_2_ and 47.5 % N_2_O. Transgenic mice were held at the animal facility of the CIPMM and fed *ad libitum*. All animals were held at a twelve-hour light/dark cycle. This study was carried out in strict accordance with the European and German guidelines for the welfare of experimental animals. Animal experiments were approved by the Saarland state’s “Landesamt für Gesundheit und Verbraucherschutz” animal licence number 36/2016.

### Golgi staining

2-month-old anesthetized mice were dislocated prior to brain dissection and 100 µm thick coronal brain sections were cut on a vibratome in the hippocampal region. The dendritic spines of hippocampal neurons from the CA1 *stratum radiatum* were visualized by the Golgi impregnation technique. For this, we used the FD Rapid GolgiStain kit (FD NeuroTechnologies). Brain sections were immersed in equal volumes of solutions A and B for 7 d and impregnated with solution C for 48 h at 4°C. Then, the sections were washed twice in double-distilled water and incubated for 10 min in a mixture solution D and solution E in double-distilled water(1:1:2). Sections were washed twice, dehydrated with increasing concentrations of ethanol, and mounted with epoxy resin (Fluka). Stacks of bright-field images with 0.3 µm spacing were acquired with a Zeiss Axioskop 50 microscope with 63x oil objective (NA 1.4; Plan-Apochromat) coupled to a CCD camera (CoolSnap ES; Roper Scientific) operated by Metaview software (Molecular Devices). Images were analysed with ImageJ. The number of dendritic spines (>1 µm protrusion) along portion of dentrite of 100 µm was counted with ImageJ.

### Transmission electron microscopy of the CA1 hippocampus

2-month-old mice mice were anesthetized and intracardially perfused with phosphate-buffered 0.9% NaCl (PBS), followed by 0.1 M phosphate buffered 4% paraformaldehyde, pH 7.4, supplemented with 0.05% glutaraldehyde (Sigma). The brains were carefully removed, postfixed for 4 h in the same fixative and 60 µm sections were cut with a vibratome. After several washes in PBS, the sections were postfixed in 1% glutaraldehyde in the same buffer for 10 min and processed for EM. This included treatment with osmium tetroxide (1% in 0.1 M PB), block staining with uranyl acetate, dehydration through a graded series of ethanol, and flat embedding on glass slides in Durcupan (Fluka) resin. Regions of interest were cut at 70 nm on an ultramicrotome (Reichert Ultracut E; Leica) and collected on one-slot copper grids. Staining was performed on drops of 1% aqueous uranyl acetate, followed by Reynolds’s lead citrate. EM images were acquired in a JEOL-1200 electron microscope with a digital camera (Veleta, SIS; Olympus) and analysed with ImageJ. Twenty images per animal from 3 animals per genotype were used for quantification. The number of synapses per µm^2^ was calculated. A synapse was considered if it met 3 criteria: a presynaptic bouton filled with at least 10 synaptic vesicles (1) juxtaposed to the head of a dendritic spine with a clearly visible PSD (2) and the presence of the neck in the section (3). Number of synaptic vesicles and areas of presynaptic boutons were quantified in each synapse using the free-shape tool and the cell counter plugins of ImageJ. We used the straight tool of ImageJ to measure the lengths of PSDs, and head and neck diameters. Note that the head diameter was taken parallel to the PSD and the neck diameter was perpendicular to the neck membranes.

### Electrophysiological recordings in CA1

#### Field recordings

Alix wt and ko brain slices were prepared from 2-month-old C57BL/6 wt and Alix ko mice. The brains were removed quickly and 350 μm-thick sagittal slices containing both cortex and hippocampus were cut in ice-cold sucrose solution (2.5 mM KCl, 1.25 mM NaH_2_PO_4_, 10 mM MgSO_4_, 0.5 mM CaCl_2_, 26 mM NaHCO3, 234 mM sucrose, 11 mM glucose, saturated with 95% O_2_ and 5% CO_2_) with a Leica VT1200 blade microtome (Leica Microsystemes, Nanterre, France). After cutting, hippocampi were extracted from the slice and transferred to oxygenated Artificial Cerebro-Spinal Fluid (ACSF: 119 mM NaCl, 2.5 mM KCl, 1.25 mM NaH_2_PO_4_, 1.3 mM MgSO_4_, 2.5 mM CaCl_2_, 26 mM NaHCO_3_, 11 mM glucose) at 37± 1°C for 30 min and then kept at room temperature for at least 1 h before recordings. Each slice was individually transferred to a submersion-type recording chamber and continuously superfused (2 ml/min) with oxygenated ACSF. Extracellular recordings were obtained at 28°C from the apical dendritic layers of the hippocampal CA1 area, using glass micropipettes filled with ACSF. Field excitatory postsynaptic potentials (fEPSPs) were evoked by the electrical stimulation of Schaffer collaterals afferent to CA1. The magnitude of the fEPSPs was determined by measuring their slope. Signals were acquired using a double EPC 10 Amplifier (HEKA Elektronik Dr. Schulze GmbH, Germany) and analysed with Patchmaster software (HEKA Elektronik Dr. Schulze GmbH, Germany). For the induction of long-term potentiation, test stimuli were delivered once every 15 s. Stimulus intensities were adjusted to produce 40-50% of the maximal response. A stable baseline was recorded for at least 15 min. LTP was induced by high frequency stimulation (4 trains delivered at 100 Hz with 5 min between each train). Average value of fEPSP slope was expressed as a percentage of the baseline response.

#### Patch-clamp recordings

Alix^fl/fl^ mice (Laporte et al., 2017) with heterozygous cre expression (Emx1^IREScre^ (Emx^tm1(cre)Krj^, MGI: 2684610)(Gorski et al., 2002) as well as control mice (Alix^fl/fl^ x Emx^wt^) were decapitated under anaesthesia and brains were removed from the skull and immersed in an ice-cold, oxygenated (5% CO_2_/95% O_2_, pH 7.4) slice preparation solution containing (in mM) 87 NaCl, 3 KCl, 25 NaHCO_3_, 1.25 NaH_2_PO_4_, 3 MgCl_2_, 0.5 CaCl_2_, 75 sucrose and 25 glucose. Coronal slices of 300 µm thickness were prepared with a vibratome (Leica VT 1200S, Nussloch, Germany) and transferred to a nylon basket slice holder for incubation in ACSF containing (in mM) 126 NaCl, 3 KCl, 25 NaHCO_3_, 15 glucose, 1.2 NaH_2_PO_4_, 2 CaCl_2_, and 2 MgCl_2_ at 32 °C for 30min. Subsequently, slices were removed from the water bath and kept at room temperature with continuous oxygenation prior to use.

Slices were transferred to the recording chamber continuously perfused with oxygenated ACSF containing (in mM) 1 MgCl_2_ and 2.5 CaCl_2_ (2-5 mL/min). During recordings, 50 μM strychnine and 50 μM picrotoxin were added to block inhibitory synaptic transmission. Pyramidal neurons were identified morphologically using the recording microscope (Axioskop 2 FS mot, Zeiss, Jena, Germany) with a 40x water immersion objective. Images were detected with a QuantEM 512SC camera (Photometrics, Tucson, USA). Whole-cell membrane currents were recorded by an EPC 10 USB amplifier (HEKA, Lambrecht, Germany), low pass filtered at 3 kHz and data acquisition was controlled by Patchmaster software (HEKA). The resistance of patch pipettes (7–9 ΩM) were fabricated from borosilicate capillaries (OD: 1.5 mm; Sutter, USA) using a Micropipette Puller (Model P-97, Sutter Instrument Co., CA). Patch pipettes were filled with a solution containing (in mM) 125 cesium gluconate, 20 tetraethylammonium (TEA), 2 MgCl_2_, 0.5 CaCl_2_, 1 EGTA, 10 HEPES and 5 Na_2_ATP (pH 7.2). The holding potential in voltage-clamp mode was at -70 mV. Spontaneous excitatory postsynaptic currents (sEPSC) were recorded under holding potential for 40 seconds in 5 minutes later after whole-cell configuration. Here only <=-10 pA sEPSC was analyzed. Excitatory postsynaptic currents (EPSCs) of pyramidal neurons in the hippocampal CA1 were evoked by stimulating Schaffer collaterals of CA3 neurons with a concentric bipolar microelectrode (MicroProbes, USA). Stimulus duration was 200 µs. To estimate short-term plasticity, 25 trains of stimuli were applied to induce synaptic depression, and the depressing trains were repeated at 10 Hz with an interval of 20 s to allow for recovery from synaptic depression. To investigate recovery from depression, stimulation with gradually increased interval was applied. The stimulation threshold was applied from 30 µA to 80 µA. All the experiments were conducted at room temperature (RT) (22 – 24 °C).

Data generated by Patch Master were loaded into Matlab (Mathworks, MA, USA) with a module adapted from sigTOOL (Lidierth, 2009). EPSC traces from the same cells were manually checked and pooled. The average of EPSC traces from each cell was used to analysis. The readily releasable pool (RRP) size was estimated by using train-extrapolation method (Neher, 2015; Thanawala and Regehr, 2016).The cumulative sum of the peak EPSC amplitudes was plotted against the stimulation numbers. The last 10 data points were linearly fitted. The RRP size was calculated by back-extrapolating the fitted line to the y-axis (stimulation ID = 0). The release probability was calculated by *p* = Amplitude^1^ / RRP_train_. The fitted slop represents the replenishment rate (Körber et al., 2015). The recovery was estimated with the decay constant (τ) as 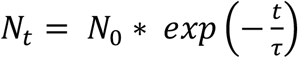.

Data analysis was performed using routines that were custom written in Matlab.

### EEG telemetry and unilateral intracortical kainate injection

We took advantage of the unilateral intracortical kainate injection model for human temporal lobe epilepsy. Telemetric EEG transmitter implantation, kainate injection and data analysis was adapted from Bedner and colleagues (Bedner et al., 2015).

Controls and alix cko were implanted with telemetric EEG transmitters (DSI PhysioTel® ETA-F10, Harvard Bioscences, Inc. Holliston, Massachusetts, USA). The animals were placed in a stereotaxic frame (Robot stereotaxic, Neurostar, Tübingen, Germany) for implantation of depth electrodes at 3.4 mm posterior to bregma and bilaterally 1.6 mm from the sagittal suture. After post-surgical care and recovery, mice were again placed in the stereotaxic frame and injected with 70 nl of a 20 mM solution of kainate (Tocris, Wiesbaden-Nordenstadt, Germany) in 0.9% NaCl, above the right dorsal hippocampus (1.9 mm posterior to bregma, 1.5 mm from sagittal suture and 1.3 mm from skull surface). Kainate was injected at a rate of 70 nl/ min with a 10 µl Nanofil syringe (34 GA blunt needle, World Precision Instruments, Sarasota, FL, USA). The syringe was kept in place for 2 min after the injection was completed to avoid liquid reflux.

Cages were placed on individual radio receiving plates (DSI PhysioTel® RPC-1, Data Sciences International, St. Paul, USA), which record EEG signals and sent them, together with the video recording (MediaRecorder Software, Noldus Information Technology, Wageningen, Netherlands), to an input exchange matrix (DSI PhysioTel® Matrix 2.0 (MX2), Ponemah software, DSI, Data Sciences International, St. Paul, USA). The animals were monitored for at least 20 h post kainate injection. In our model the mortality rate associated to *status epilepticus* is less than 5% in more than 50 mice with different genetic backgrounds over the last 12 months.

EEG traces were analysed with the Neuroscore software (Version 3.3.1., Data Sciences International, St. Paul, USA). Electrographic seizures were detected with the spike detection protocol. Subsequently, an additional manual screen was employed to remove artifacts that were eventually picked up. Seizures were characterized by high frequency spiking and ceased with a postictal depression (flattening of EEG). Seizure detection was complemented by synchronized video monitoring. Electrographic seizures were associated with behavioural analogues of Racine stages II-V (Racine, 1972). The total duration of *status epilepticus* was defined from the first electrographic seizure to the first seizure free period lasting 1 h.

For immunohistochemitry analysis, 40 µm free-floating vibratome sections were incubated for one hour in blocking buffer (5 % HS, 0.3 % Triton X in 1x PBS) at room temperature. Sections were incubated with primary antibodies (see antibodies section) diluted in blocking solution, overnight at 4°C. Secondary antibodies and DAPI (0.025 μg/ml, Sigma) were diluted in blocking buffer and incubated for 2 h at room temperature.

### Cell culture and transfection

Cortical and hippocampal neurons from rat E18 embryos and P0 Alix wt and ko mice were prepared as previously described (Faure et al., 2006). Briefly, cortices and hippocampi were dissected from E18 rat embryos or P0 mouse pups, treated with trypsin, and mechanically dissociated. Dissociated cells were seeded at a density of 5 × 10^4^/cm^2^ in 100 mm dishes for cortical neurons and 1.5 × 10^4^/cm^2^ onto acid-washed coverslips (either 14 mm diameter or 25 mm; Marienfeld) in either 4-well plates or P35 dishes (ThermoScientific) precoated for 4 h with 50 μg/ml poly-D-lysine (Sigma). Neurons were plated in DMEM containing 10 unit/mL penicillin, 10 μg/mL streptomycin,0.5 mM L-glutamine and 10% of inactivated horse serum (Invitrogen) Neurons were maintained in water-saturated 95% air/5% CO_2_ at 37°C. The seeding medium was replaced after 20 h with serum-free neuronal culture medium (Neurobasal medium containing 2% B27 supplement, 10 unit/mL penicillin, 10 μg/mL streptomycin, and 0.5 mM L-glutamine; Invitrogen). Neurons were transfected at 10 DIV as previously described (Chassefeyre et al., 2015). Briefly, for each P35 dish, 2 μg plasmid DNA, 250 mM CaCl_2_ were mixed with an equal volume of 2x BES-buffered saline and left to precipitate for 20 min at room temperature. Neurons were placed in transfection medium (Minimum Essential Medium containing 0.22% NaHCO_3_, 20 mM D-glucose and 0.5 mM L-glutamine) supplemented with 2% B27, before the DNA precipitate was added. They were then incubated for 1.5 h at 37°C and 5% CO_2_. Neurons were then washed by being placed in transfection medium (pre-warmed to 37°C in 10% CO_2_) for 20 min at 37°C and 5% CO_2_. Finally, they were transferred back into their conditioned medium.

Primary cultures of cerebellar granule neurons (CGN) were prepared from 6-day-old C57BL/6 Alix wt and Alix ko pups as described previously (Trioulier et al., 2004), with some modifications. The cerebella were removed, cleared of their meninges, and cut into 1-mm pieces. They were then incubated at 37°C for 10 min in 0.25% trypsin-EDTA and DNAse (1500 U/mL). Trypsin was inactivated and cells were dissociated in culture medium (DMEM containing 10% fetal bovine serum, 2 mM L-glutamine, 25 mM KCl, 10 mM HEPES and 10 unit/ml penicillin, 10 µg/ml streptomycin). After filtration on 70 µm cell strainers, neurons were plated at 5.10^5^ cell/cm^2^ onto poly-D-lysine (10 µg/ml, Sigma) precoated coverslips. Cytosine-β-D-arabinoside (10 µM, Sigma) was added after 1 day in vitro (DIV), to prevent the growth of non-neuronal cells, until 8 DIV when neurons were used for activity dependent bulk endocytosis experiment (see below).

### Multi-electrode array

Dissociated hippocampal neurons were resuspended in neuronal medium and plated at a 10^6^ cells/cm^2^ on poly-L-lysine-coated multi-electrode arrays comprising 59 extracellular recording electrodes and one reference electrode (MEA-60 Multichannel Systems, MCS). Electrodes were 30 µm in diameter and separated by a distance of 200 µm. MEA-60 plates were connected to a 60-channel data acquisition system (USB-MEA64) and associated amplifier (MEA1060-Inv-BC) powered by a PS40W power supply system (MCS, Germany). The recording system was then placed in a humidified incubation chamber at 37°C and 5% CO_2_. Neurons were left for 5 min to equilibrate before recording. Basal spontaneous activity was recorded for 3 min prior addition of drugs. Bicuculline (100 µM) and 4-aminopyridine (5 mM) were then added and neuronal activity recorded for another 10 min. Signals were recorded with a 1100 gain, sampled at 20 kHz and analyzed with MC Rack software (MCS, Germany). Raw signals were first filtered with a Butterworth band pass filter (2nd order) between 200 and 800 Hz to remove electrical noise and low frequency signals. Spike detection analysis was performed on filtered signals using a threshold of 6 standard deviations to mark out action potentials. Burst events were identified as a minimum of 5 consecutive spikes with an interspike interval lower than 100 ms. The minimum interval time between two bursts was fixed at 1000 ms. Representative traces were exported using custom-made Matlab functions (Matlab 2014b).

### Live imaging of synaptophysin-pHluorin upon electrical stimulation

Hippocampal neurons from Alix wt and ko mice were transfected with synaptophysin-pHluorin (Syp-pH) at 6 DIV. The recycling of synaptic vesicles was imaged in a buffer solution containing 120 mM NaCl, 5 mM KCl, 2 mM CaCl_2_, 2 mM MgCl_2_, 5 mM glucose, 10 mM HEPES adjusted to pH 7.4 and 270 mOsm/l. Experiments were carried out at 34°C. Neurons were stimulated by electric field stimulation (platinum electrodes, 10 mm spacing, 1 ms pulses of 50 mA and alternating polarity at 5-40 Hz) applied by a constant current stimulus isolator (SIU-102, Warner Instruments). The presence of 10 µM 6-cyano-7-nitroquinoxaline-2,3-dione (CNQX) and 50 µM D,L-2-amino-5-phosphonovaleric acid (AP5) prevented recurrent activity. Experiments were performed on an inverted microscope (IX83, Olympus) equipped with an Apochromat N oil 100× objective (NA 1.49). Images were acquired with an electron multiplying charge coupled device camera (QuantEM:512SC; Roper Scientific) controlled by MetaVue7.1 (Roper Scientific). Samples were illuminated by a 473-nm laser (Cobolt). Emitted fluorescence was detected after passing a 525/50 nm filter (Chroma Technology Corp.). Time-lapse images were acquired at 1 or 2 Hz with integration times from 50 to 100 ms.

Image analysis was performed with custom macros in Igor Pro (Wavemetrics) using an automated detection algorithm as described previously (Martineau et al., 2017). The image from the time series showing maximum response during stimulation was subjected to an “à trous” wavelet transformation. All identified masks and calculated time courses were visually inspected for correspondence to individual functional boutons. The intensity values were normalized to the ten frames before stimulation.

### Live Fluorescence Imaging of Alix and endophilin recruitment to synapses

For live imaging of protein recruitment, cultured hippocampal neurons were co-transfected with Alix-mCherry or endophilin-A2-mCherry together with Syp-pH expression vectors 2 to 4 days prior to imaging. All live imaging experiments were performed at 37°C and images were acquired using a spinning disk confocal microscope (AxioObserver Z1) with a 63x oil objective (NA 1.46, Zeiss). Transfected neurons were placed in basal medium for 10 min and then mounted in an imaging chamber (POC-R2 Cell cultivation system, Zeiss). After 2 min in basal medium, a 5x bicuculline/4AP solution was added (final concentration, 100 μM and 5 mM, respectively) for stimulation which lasted 5 min and the chamber was perfused with basal medium at 3 ml/min for washing for 5min. ROI were drawn on ‘presynapses’ defined by spots of synaptophysin-pHluorin that increased during stimulation. Fluorescence values were measured in these ROI and then normalised to the initial fluorescence values (fluorescence values prior to stimulation) using ImageJ.

### Immunofluorescence

Cultured hippocampal neurons were fixed for 20 min at room temperature in phosphate-buffered 4% paraformaldehyde supplemented with 4% sucrose. After three washes in PBS, cells were permeabilized and blocked in PBS containing 0.3% Triton X-100 and 3% BSA for 15 min at room temperature. Coverslips were incubated for 1–2 h at room temperature with primary antibodies (see ‘antibodies’ section) diluted in the blocking solution. After washing in PBS, cells were incubated for 1 h with secondary antibodies conjugated to Alexa Fluor 488, Alexa Fluor 594, or Cyanine 5 (Cy5), diluted in the blocking solution. Coverslips were rinsed and mounted in Mowiol. Images were acquired on a Leica SPE microscope using a 40x dry objective (NA 0.75, Leica) or a 100x oil immersion objective (NA 1.4, Leica) at 488 nm, 532 nm or 635 nm.

#### Synapse density

Cultured hippocampal neurons were fixed and stained with antibodies at 14-16 DIV as described above. Presynaptic boutons (synapsin-1-positive) and postsynaptic terminals (PSD95-positive) were selected using the Spot Detector plugin in Icy software (de Chaumont et al., 2012) (wavelet detection with size filtering between 0.4 μm and 2 μm in diameter) on max image projections. Synapses were defined as spots of colocalization between the detected presynaptic and postsynaptic terminals that were within 3 μm of each other. Synapses were counted using the Colocalization Studio in ICY software (Lagache et al., 2018)

#### Protein recruitment assays

Cultured hippocampal neurons were transfected with expression vectors 24∼48 h prior to stimulation. The transfected neurons were incubated in basal medium (150 mM NaCl, 5 mM KCl, 1.3 mM CaCl_2_, 10 mM HEPES and 33 mM D-glucose at pH 7.4) for 10 min prior to stimulation. Coverslips were either treated with basal medium alone or bicuculline/4AP solution (basal medium supplemented with 100 μM bicuculline and 5 mM 4-aminopyridine) for 5 min at 37°C and 5% CO_2_. Coverslips were fixed and stained with antibodies against synapsin-1 and PSD95 (see ‘antibodies’ section). Images were acquired on a Leica SPE microscope using a 100x oil immersion objective (NA 1.4, Leica). The acquired images were analysed using ImageJ. Regions of interest (ROI) were drawn on areas of axons which colocalize with anti-synapsin-1 spots, but not with anti-PSD95 spots to ensure presynaptic measurement. Fluorescence intensities were then measured in these ROI and normalised to fluorescence intensities measured on other regions of axons to give the relative fluorescence intensity at presynaptic boutons.

### Calcium increase imaging upon neuronal stimulation

Hippocampal neurons cultured on 24-well plates were placed in basal medium and incubated with 1 μM Fluo-4-AM for 1 h at 37°C. After washing, neurons were left in 250 μl basal medium for 5 min. Plates were then transferred to a Pherastar automatic plate reader (BMG Labtech, Germany) set to 37 °C, to record fluorescence intensity at 0.1 Hz during 10min, for each well, and with fluorescence measurement settings as described on http://www.bmglabtech.com/media/35216/1043854.pdf. Neurons were stimulated with 50 μl 6x bicuculline/4AP solution (final concentration, 100 μM and 5 mM, respectively) after 5min with automatic injection.

### Dextran uptake upon neuronal stimulation

The protocol for dextran uptake was adapted from (Clayton and Cousin, 2009b). 15-17 DIV hippocampal neurons were stimulated with bicuculline/4AP solution for 5 min, at 37°C and 5% CO_2_, in the presence of 10 kDa tetramethylrhodamine-dextran (50 μM). Coverslips were immediately washed several times in washing solution (basal medium supplemented with 0.2% BSA and warmed to 37°C) to remove excess dextran. For ADBE inhibition, neurons were stimulated in the presence of 2 μM GSK3 inhibitor (CT99021, Tocris) and placed in fresh basal medium containing 2 μM GSK3 inhibitor for 10 min at 37°C and 5% CO_2_. Neurons were then fixed as previously described and imaged with a Leica SPE microscope using a 40x dry immersion objective (NA 0.75, Leica). Analysis was performed on ImageJ. The number of fluorescent spots was counted in a defined field of view (130μm x 130 μm) in thresholding analysis with a diameter limit between 300 nm and 2 μm (resolution limit for the microscope and maximum size of a nerve terminal).

For rescue experiments, cultured hippocampal neurons were transfected 2 to 4 d prior to the day of experiment with the following constructs :Alix wt, Alix I212D (AlixΔCHMP4B), AlixΔPGY (AlixΔALG-2), Alix R757E (AlixΔendo). Dextran uptake assay was performed as described above. Images were acquired on a Leica SPE microscope using a 40x oil immersion objective (NA 1.25, Leica) at 488 nm and 532 mm excitation. The analysis was performed on ICY software. ROI were generated closely around axons by using ‘thresholder’. Then dextran spots within these ROI were counted by using ‘spot detector’ with size limit between 300 nm and 2 μm. The length of axon per field of view was estimated by manually drawing ‘Polyline type ROI’ over the axon images. The number of dextran spots per μm of axon was calculated and expressed as ratio to control values.

### EM examination of ADBE in culture neurons

Analysis of ADBE from cerebellar granule neurons was performed as described previously with some modifications (Cheung et al., 2010). 8 DIV CGN were incubated in hyperpolarizing medium (170 mM NaCl, 3.5 mM KCl, 0.4 mM KH_2_PO_4_, 20 mM TES, 5 mM NaHCO_3_, 5 mM D-glucose, 1.2 mM Na_2_SO_4_, 1.2 mM MgCl_2_, 1.3 mM CaCl_2_, pH7.4) for 10 min prior to stimulation. Neurons were then incubated for 2 min with 10 mg/ml HRP (Sigma P8250) in either the hyperpolarizing medium or a high-potassic solution containing 50 mM KCl (123.5 mM NaCl, 50 mM KCl, 0.4 mM KH_2_PO_4_, 20 mM TES, 5 mM NaHCO_3_, 5 mM D-glucose, 1.2 mM Na_2_SO_4_, 1.2 mM MgCl_2_, 1.3 mM CaCl_2_, pH7.4) before rapid washing in PBS and fixation in PBS-glutaraldehyde 2% for 30 min. After three washes in Tris buffer 100 mM, endocytosed HRP was revealed by incubation in Tris 100 mM containing 0.1% diaminobenzidine and 0.2% H_2_O_2_. The cultures were then post-fixed in 1% osmium tetroxide, dehydrated and embedded in Epon. Synapses were photographed with a JEOL-1200 electron microscope. Quantification was performed as follows: HRP-positive structures were quantified per synapse and classified as synaptic vesicles when the diameter was less than 100 nm and as bulk endosome when the diameter was more than 100 nm.

For focused ion beam scanning electron microscopy (FIB-SEM), the block was mounted on a pin, coated with gold, and inserted into the chamber of a HELIOS 660 Nanolab DualBeam SEM/FIB microscope (FEI Co., Eindhoven, The Netherlands). Region of interest (ROI) was prepared using focused ion beam (FIB) and ROI set to be approximatively 15 µm wide. During the acquisition process, the image acquisition parameters of the electron beam were 2 kV and 0.4 nA and the thickness of the FIB slice between each image acquisition was 10 nm. The segmentation of the synapse and ABDE was done with Amira software (Thermo Fisher Scientist) and the movie obtained using Imaris (Oxford Instruments).

### Synaptosomal preparation from cortical neurons

Synaptosome-enriched membranes from 15 DIV cortical neurons were prepared as described previously with some modifications (Frandemiche et al., 2014). Briefly, cultured neurons were stimulated for 15 min with a mixture of bicuculline/4AP (50 µM/2.5 mM). After a wash in HBSS, neurons were homogenized by passing 15-20 times through 0.25G needle in cold buffer containing 0.32 M sucrose, 10 mM HEPES, 15mM NaF, 15mM β-glycerophosphate and protease inhibitors (Roche), pH 7.4. Samples were maintained at 4°C during all steps of the experiment. Homogenates were cleared at 1000g for 10 min to remove nuclei and large debris. The resulting supernatants were spun down at 12,000 g for 20 min to obtain a crude membrane fraction and washed twice in HEPES buffer 4 mM containing 1 mM EDTA, 15 mM NaF, 15 mM β-glycerophosphate and protease inhibitors (Roche), pH 7.4. The resulting pellet was solubilized in 0.5% Triton X-100, 20 mM HEPES, 100 mM NaCl, 15mM NaF, 15 mM β-glycerophosphate, pH 7.2, containing protease inhibitors (Roche) for 20 min at 4°C with mild agitation and analysed by Western blot.

### Western blot

Cells lysates were resuspended in Laemmli buffer and resolved by SDS-PAGE in 10% polyacrylamide gels. Proteins were electro-transferred onto PVDF membranes that were then blocked for 30 min in TBS containing 0.1% Tween 20 and 5% dry milk and incubated for 1 h to overnight with primary antibodies diluted in the blocking solution. After washes in TBS– Tween, the membranes were further incubated for 1 h with secondary antibodies coupled to HRP, washed as before and incubated with luminescence-generating HRP substrate. Bound antibodies were revealed by luminography on film.

### Statistical analysis

The comparison of two groups was performed using two-sided Student’s t-test or its non-parametric correspondent, the Mann-Whitney test, if normality was rejected (Shapiro-Wilks test). The comparisons of more than two groups were made using one or two way ANOVAs followed by post-hoc tests (Holm Sidak’s or Tukey’s HSD) to identify all the significant group differences. N indicates independent biological replicates. The graphs with error bars indicate 1 SEM (+/-) except for figure 5 where we used box plots showing distribution of medians (whiskers = min and max values), figures 7 and supplementary figure 6 where median and interquartile range (IQR) were represented. The significance level is denoted as usual (*p<0.05, **p<0.01, ***p<0.001). All the statistical analyses were performed using Prism7 (Graphpad version 7.0a, April 2, 2016).

## Results

### Alix is recruited to synapses upon synaptic activation

We used dissociated cortical neuron cultures to decipher the role of Alix at synapses. Western blot analysis during *in vitro* differentiation revealed that Alix expression strongly increases during synaptogenesis as indicated by the parallel rise in postsynaptic density protein 95 (PSD95) expression (**Figure 1A**). Moreover, synaptosome-enriched membranes prepared from mature cortical neurons contained Alix (**Figure 1B**). This synaptic pool increased when neuron cultures were incubated 15 min with the GABA_A_ receptor antagonist bicuculline, together with a weak potassium channel blocker 4-aminopyridine (4AP)(**Figure 1B**), a treatment known to increase the frequency of action potential bursts (**Supplementary figure 3C**) and thereby induce sustained intracellular calcium elevation (**Supplementary figure 1A**) (Hardingham et al., 2001; Hardingham et al., 2002). Therefore, this observation suggests that Alix tends to concentrate at synapses undergoing prolonged stimulations. To further confirm this, we used live imaging to follow mCherry-Alix relocalization to synapses in mature hippocampal neurons. While the fluorescent signal was homogeneously distributed throughout the entire neuronal cytoplasm in resting conditions (**Figure 1C**, t= 0 min), it almost doubled in discrete spots within neuronal processes during synaptic activation (**Figure 1C**, t=2m20s arrowheads). These spots correspond to active presynaptic boutons as they were labelled with Synaptophysin-pHluorin (Syp-pH), which becomes fluorescent upon exocytosis at synapses (Granseth et al., 2006) (arrowheads **Figure 1C**; **Movie 1**). Alix recruitment to these sites is concomitant with the activation of glutamatergic synapses, being detectable a few seconds after addition of bicuculline/4AP to the culture medium and decreasing soon after washing the cells (**Figure 1 C, D**). Alix relocalization was also seen with Alix-YFP which concentrated during bicuculline/4AP treatment at sites confirmed to be at axonal boutons by synapsin-1 immunolabelling (**Figure 1E, F**). Anti-PSD95 immunostaining showed that Alix-YFP positive spots were juxtaposed to, but did not overlap with the postsynaptic density marker (**Figure 1G**). Accordingly, measuring Alix-YFP fluorescence intensities in dendritic spines before and during stimulation showed no accumulation of Alix at the post-synaptic level (**Supplementary figure 1B, C**), further demonstrating that synaptic activation leads to Alix recruitment only at presynaptic parts.

**Figure 1.**
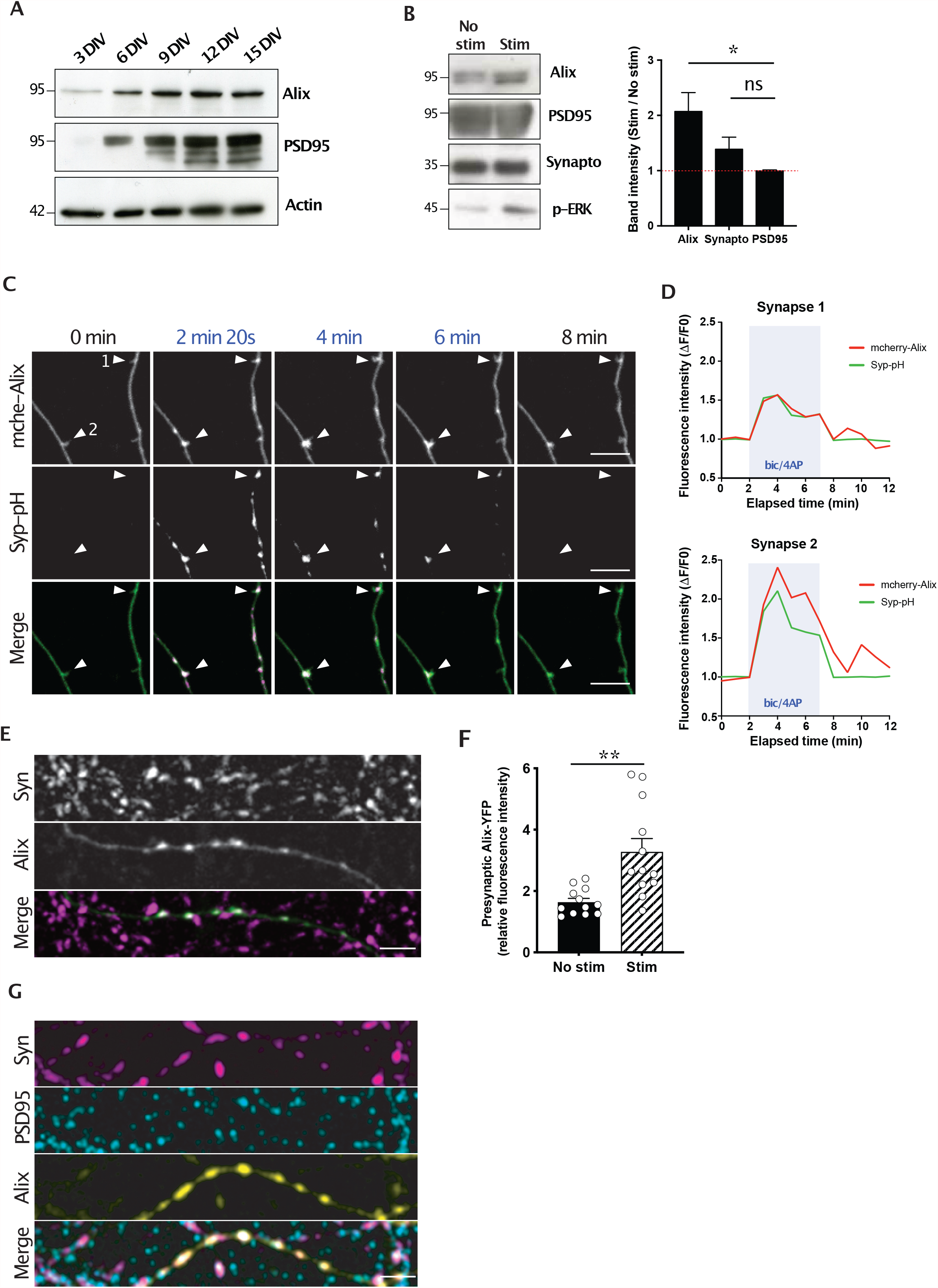
Alix is recruited presynaptically during synaptic activation. **A**) Western blot analysis of cortical neurons cultured for 3 to 15 DIV demonstrates the increase in Alix expression correlating with synaptogenesis as illustrated by the increase in PSD95 expression. **B**) Western blot analysis of the increase of Alix in synaptosome-enriched neuronal membranes upon neuronal stimulation by bicuculline/4AP. Synaptophysin and PSD95 were used as pre- and postsynaptic markers, respectively. The phosphorylated form of ERK (p-ERK) assessed the efficiency of the stimulation. **C**) Images from time-lapse video microscopy of 15 DIV hippocampal neurons expressing both mCherry-Alix (mche-Alix) and synaptophysin-pHluorin (Syp-pH) stimulated between 2 and 7 minutes with bicuculline/4AP. White arrowheads indicate presynaptic boutons where Alix is recruited during stimulation. Scale bar: 10 μm. **D**) Profiles of mcherry-Alix and Syp-pH fluorescent increase at presynaptic boutons 1 and 2 of panel **C**. Blue box indicates the bicuculline/4AP stimulation. **E**) 15 DIV hippocampal neurons expressing Alix-YFP (green) stimulated for 5 min before fixation and stained with anti-synapsin-1 antibody (Syn, magenta). Scale bar: 5 μm. **F**) Graph shows the presynaptic increase in Alix-YFP upon stimulation. Presynaptic Alix-YFP corresponds to the ratio of YFP fluorescence between synapsin-positive and - negative axonal regions. **G**) 15 DIV hippocampal neurons expressing Alix-YFP (green) stimulated for 5 min before fixation and stained with anti-synapsin-1 antibody (Syn, magenta) and anti-PSD95 (post-synaptic, cyan) showing the selective recruitment of Alix to presynpase. Scale bar: 5 μm. <u>Average +/-SEM, N, statistical analysis:</u> **B)** 2.07+/-0.34; 1.38+/-0.21; 1.00+/-0.01 for Alix, Synaptophysin and PSD95 respectively. N=4 independent experiments, Alix versus PSD95, p=0.0187, one-way ANOVA. **F**) 1.63+/-0.12; 3.27+/-0.44 for No Stim and Stim respectively. N=12 neurons per condition from 4 independent experiments, p=0.0017, Unpaired t-test.

### Recruitment of Alix partners, ALG-2 and endophilin-A, at activated synapses

In non-neuronal cells, Alix recruitment to plasma membrane depends on calcium binding to ALG-2, a penta-EF-hand containing protein, which thereafter interacts and activates Alix (Scheffer et al., 2014). In neurons, action potential depolarization induces massive and transient calcium accumulation in the bouton, which triggers fusion of SV with the plasma membrane. Adding the intra-cellular calcium chelators BAPTA-AM or EGTA-AM completely abolished Alix-YFP recruitment (**Supplementary figure 2**). Interestingly, GFP-ALG-2 concentrated at presynaptic boutons upon synaptic activation, in contrast to an ALG-2 point-mutant unable to bind calcium (ALG-2ΔCa) (Shibata et al., 2004) (**Figure 2A, B**). The use of neurons prepared from Alix ko mice (Laporte et al., 2017; Mercier et al., 2016) showed that ALG-2 recruitment does not depend on Alix, as it concentrates at stimulated synapses of Alix ko neurons at the same level than in wt neurons (**Figure 2B, C** left histograms). On the contrary, Alix recruitment to active synapses is tightly dependent on its capacity to interact with ALG-2, as a mutated version of the protein unable to interact with ALG-2 (AlixΔALG-2) (Suzuki et al., 2008; Trioulier et al., 2004) does not accumulate at presynaptic boutons upon stimulation (**Figure 2C**, right histogram).

**Figure 2.**
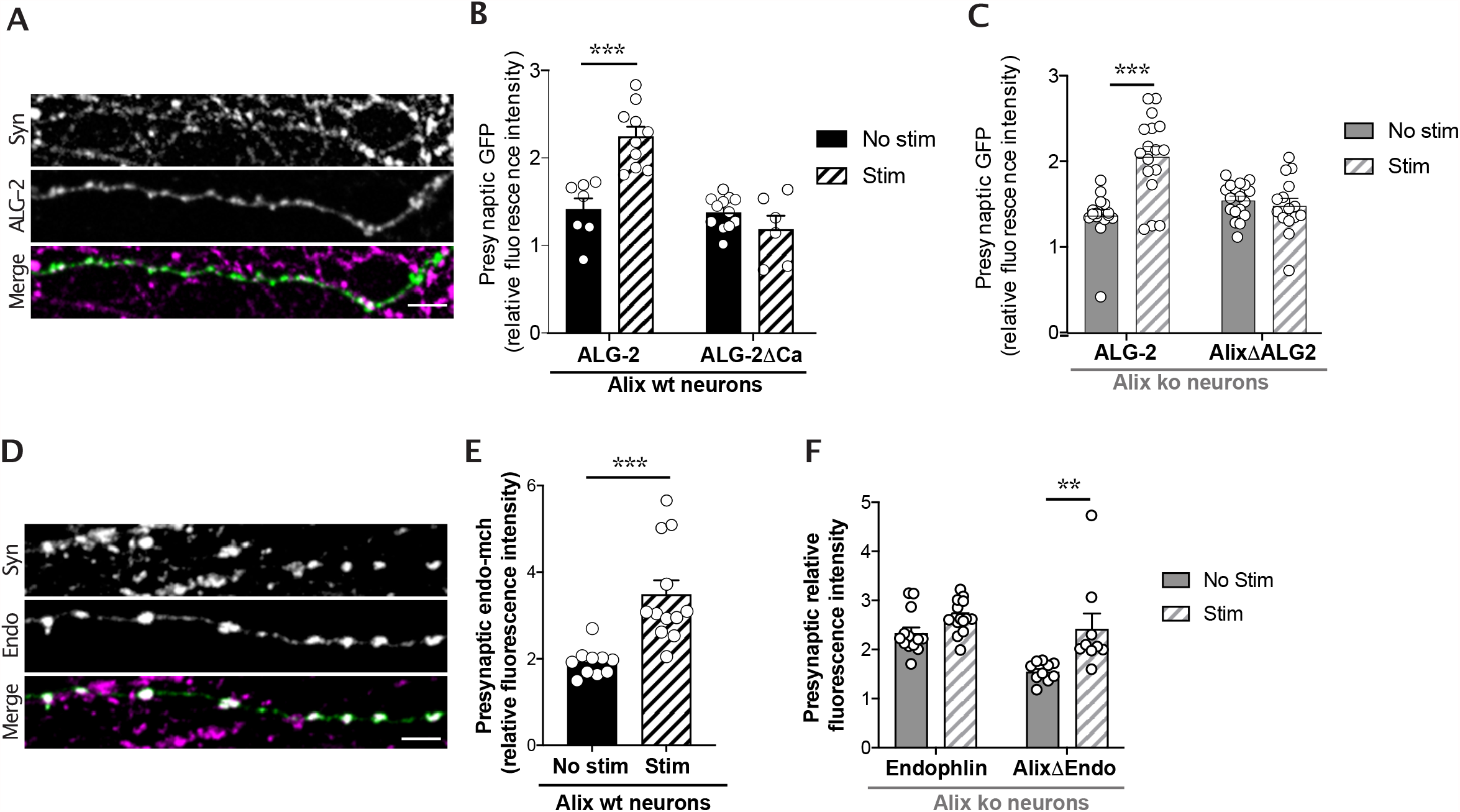
Interplay between Alix, ALG-2 and endophilin recruitments at activated synapses. All experiments made use of 15 DIV hippocampal neurons expressing the indicated constructs. Neurons were stimulated with bicuculline/4AP for 5 min before fixation and stained with anti-synapsin-1 (Syn, magenta). **A**) GFP-ALG-2 (green) is recruited presynaptically upon stimulation. Scale bar: 5 μm. **B**) Quantification of synaptic recruitments of GFP-ALG-2 and GFP-ALG-2ΔCa in Alix wt neurons shows that ALG-2 presynaptic increase depends on its capacity to bind calcium. **C**) Alix recruitment depends on its capacity to bind ALG-2 as GFP-AlixΔALG-2 is not recruited upon stimulation of Alix ko neurons. However, ALG-2 does not require Alix to be recruited as GFP-ALG-2 is recruited presynaptically in Alix ko neurons. **D, E**) Endophilin-mCherry (endo, green) concentrates at presynaptic parts of stimulated neurons. Scale bar: 5 μm. **F**) Endophilin concentration following synaptic activation requires Alix as it does not occur in Alix ko neurons. Alix recruitment does not depend on its binding to endophilins as shown with GFP-AlixΔendo in Alix ko neurons. <u>Average +/-SEM, N, statistical analysis:</u> **B)** 1.41+/-0.13; 2.24+/-0.11; 1.38+/-0.05; 1.18+/-0.15 for GFP-ALG-2 no stim, GFP-ALG-2 stim, GFP-ALG-2ΔCa no stim, GFP-ALG-2ΔCa stim respectively. N=7 and n=10 neurons for GFP-ALG-2 not stimulated and stimulated, respectively and n=12 and n=6 neurons for GFP-ALG-2ΔCa not stimulated and stimulated, respectively, from 3 experiments, p=0.0001, one-way ANOVA. **C)** 1.37+/-0.07; 2.06+/-0.11; 1.54+/-0.05; 1.48+/-0.08 for GFP-ALG-2 no stim; GFP-ALG-2 stim, GFP-AlixΔALG-2 no stim, GFP-AlixΔALG-2 stim respectively. N=17 neurons for both GFP-ALG-2 not stimulated and stimulated, and n=18 and n=15 neurons for GFP-AlixΔALG-2 not stimulated and stimulated, respectively, from 3 independent experiments, p=0.0001, one-way ANOVA. **D**,**E)** 1.92+/-0.11; 3.48+/-0.33 for no stim and stim respectively. N=10 and n=12 neurons for not stimulated and stimulated, respectively from 3 independent experiments, p=0.0001, Mann Whitney test. **F)** 2.37+/-0.12; 2.66+/-0.08; 1.56+/-0.06; 2.43+/-0.32 for endophilin-mCherry no stim, endophilin-mCherry stim, GFP-AlixΔendo no stim, GFP-AlixΔendo stim respectively. N=13 and n=15 neurons for endophilin-mCherry not stimulated and stimulated, respectively, and n=10 and n=9 neurons for GFP-AlixΔendo not stimulated and stimulated, respectively, from 3 independent experiments p=0.0031, one-way ANOVA.

**Figure 3.**
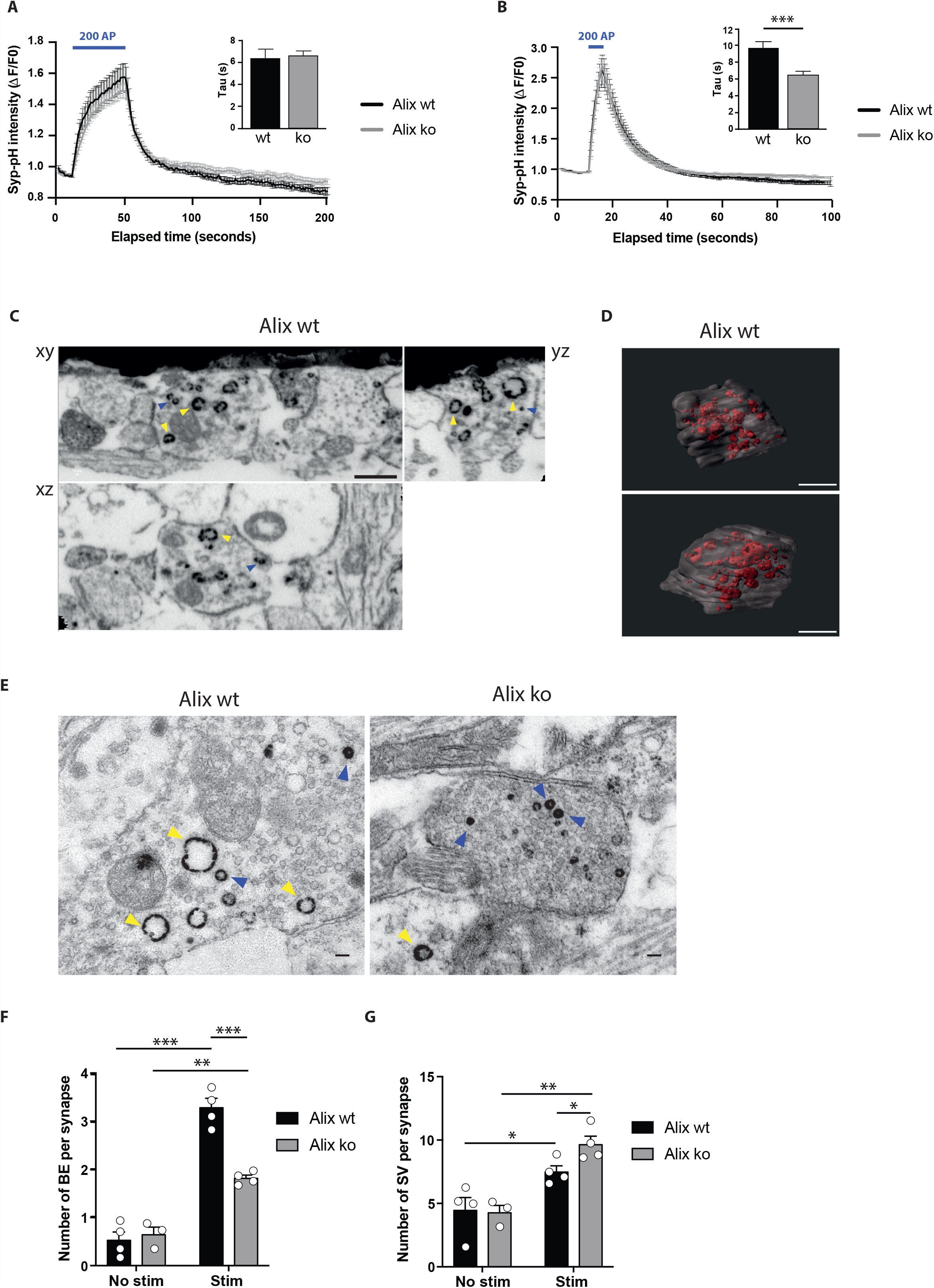
Alix is necessary for activity-dependent bulk endocytosis. **A, B**) Average traces of Syp-pH fluorescence in synaptic boutons of Alix wt and ko hippocampal neurons stimulated with 200 action potentials (AP) applied at 5 Hz (**A**) or 40 Hz (**B**). Insets show the exponential fit of fluorescence decay after stimulations in the fields imaged. **C, E**) Electron micrographs of cerebellar granule neurons stimulated in presence of free HRP to label newly-formed synaptic vesicle (blue arrowhead) and bulk endosomes (yellow arrowheads). **C**) Focused ion beam scanning electron microscopy (FIB-SEM); orthogonal views from different planes (xy, xz, yz) extracted from a stack used for the 3D reconstruction of wt presynaptic terminal shown in (**D**). Scale bar: 500nm. **D**) Two different views of the reconstructed synapse are shown where the membrane is represented in transparent gray and HRP-positive structures in red. Scale bar: 500nm. E) Transmission electron microscopy of HRP incubated CB Alix wt and Alix ko neurons. Scale bar: 100 nm. **F, G**) Quantification of the number of bulk endosomes (**F**) and synaptic vesicles (**G**) in Alix wt and ko neurons from images as shown in (E). <u>Average +/-SEM, N, statistical analysis:</u> **A, B**) N= 16-46 fields of view per condition from 4 experiments for wt and 5 for ko mice, p=0.99 (**A**) and p=0.001 (**B**), one-way ANOVA. **F, G**) 0.52+/-0.17; 0.64+/-0.15; 3.28+/-0.19; 1.82+/-0.06 for Alix wt no stim, Alix ko no stim, Alix wt stim, Alix ko stim respectively (**F**). 4.43+/-1.02; 4.26+/-0.58; 7.46+/-0.49; 9.64+/-0.67 for Alix wt no stim, Alix ko no stim, Alix wt stim, Alix ko stim respectively (**G**). N= 44 and 51 neurons from 4 independent experiments, p<0.0001 (Alix wt no stim vs Alix wt stim, **F**), p=0.0015 (Alix ko no stim vs Alix ko stim, **F**), p=0.0001 (Alix wt stim vs Alix ko stim, **F**) and p=0.0376 (Alix wt no stim vs Alix wt stim, **G**), p=0.0014 (Alix ko no stim vs Alix ko stim, **G**), p=0.0393 (Alix wt stim vs Alix ko stim, **G**), one-way ANOVA.

Among demonstrated interactors of Alix, endophilins-A are main regulators of endocytosis at synapses and impact the number of SVs (Milosevic et al., 2011; Schuske et al., 2003). We therefore tested if endophilins-A could be recruited to active synapses, similarly to Alix and ALG-2. Endophilin-A2-mCherry was mainly detected at presynaptic boutons and its fluorescence increased during bicuculline/4AP treatment (**Figure 2D, E; Movie 2**). Remarkably, no such increase could be seen in Alix ko neurons, whereas Alix deleted of its endophilin binding domain (AlixΔendo) (Chatellard-Causse et al., 2002), was still able to be recruited to synapses upon activation (**Figure 2F**). This strongly suggests that the increase in endophilin concentration during sustained synaptic activation requires Alix expression.

### Alix is required for ADBE

Altogether, our observations indicate that calcium increase at synapses undergoing sustained activity allows the transient recruitment of ALG-2, Alix and endophilin-A sequentially to presynaptic parts. Because of the demonstrated roles of Alix in clathrin-independent endocytosis, we next tested if the turnover of presynaptic vesicles might be affected by the lack of the protein by quantifying Syp-pH fluorescence during electrical stimulation. Syp-pH fluorescence increases during stimulation witnessing exocytosis. It decays thereafter reflecting SV retrieval and vesicle re-acidification (Soykan et al., 2017). As shown in **figures 3A and B**, no striking difference in the increase of fluorescence was detected between Alix ko and wt neurons stimulated at 5 or 40 Hz. However, the time constant of the exponential decay was slightly decreased in Alix ko neurons stimulated at 40 Hz, suggesting a higher rate of Syp-pH endocytosis occurring at this frequency (**Figure 3B**).

We next tested if the proteins might intervene in activity-dependent bulk endocytosis (ADBE) at synapses. Cultured cerebellar granular neurons (CGN) have been extensively used to study this process (Cheung et al., 2010). Here, neurons were depolarized with 50 mM KCl, in the presence of Horse Radish Peroxidase (HRP) that is endocytosed and fills vesicles and endosomes as they form. Examination of wt synapses by electron microscopy showed that depolarization dramatically increased the number of HRP-positive vacuoles. We verified using FIB SEM and 3D reconstruction that these vacuoles had undergone fission from the plasma membrane and could therefore be identified as bulk endosomes (**Figure 3C** yellow arrowheads; **Figure 3D, E**). Other HRP-positive vesicles having the size of neurotransmitter vesicles were also more numerous in depolarized synapses (**Figure 3C** blue arrowheads; **Figure 3D, E**). In Alix ko synapses, the number of depolarization-induced bulk endosomes was strongly reduced (**Figure 3E, F**). Consistent with the increased endocytic rate detected by Syp-pH in Alix ko neurons stimulated at 40 Hz (**Figure 3B**), the number of newly formed SV was significantly increased, suggesting a mechanism compensating for ADBE deficiency (**Figure 3E, G**).

Another way to assess for ADBE is by the use of fluorescent 10 kDa dextran, a fluid phase cargo that accumulates inside bulk endosomes but fails to label vesicles formed by clathrin-dependent mechanisms upon neuronal stimulation (Clayton and Cousin, 2009a). Hippocampal neurons incorporated dextran when incubated in presence of bicuculline/4AP (**Figure 4A, B**). A GSK3 inhibitor, known to block ADBE but no other modes of SV endocytosis (Clayton et al., 2010), completely abolished the dextran labelling of wt hippocampal neurons (**Supplementary figure 3A, B**). In contrast, Alix ko neurons failed to endocytose dextran (**Figure 4A, B**) even though bicuculline/4AP stimulation increased calcium entry (**Supplementary Figure 1A**) and neuronal activity (**Supplementary figure 3C-F**) in both Alix ko and wt neurons. Noteworthy is that even though bicuculline/4AP increased activity in both cases, the frequency of burst was significantly reduced in Alix ko neurons (**Supplementary figure 3C, D**). As already shown (Morton et al., 2015), neurons also failed to endocytose dextran when treated with calcium chelators BAPTA and EGTA (**Supplementary figure 3E**), both of which blocked Alix recruitment to presynaptic boutons (**Supplementary figure 2**). Finally, expression of GFP-Alix constructs in Alix ko neurons showed that the impairment in ADBE observed in Alix ko cells is due solely to the absence of Alix, since Alix expression fully restored the capacity of Alix ko neurons to endocytose 10 kDa dextran (**Figure 4 C, D**). Rescue was also obtained with AlixΔ4B, a mutant unable to bind the ESCRT-III protein CHMP4B. This was not the case with AlixΔALG-2 or AlixΔEndo which were not able to rescue dextran endocytosis (**Figure 4D**). Noteworthy is that AlixΔALG-2 cannot be recruited to synapses but both AlixΔendo and AlixΔ4B are. Thus, besides its requirement for ALG-2-binding to be recruited at synapses, Alix needs to interact with endophilins but not with CHMP4B-ESCRT-III in order to drive ADBE.

**Figure 4.**
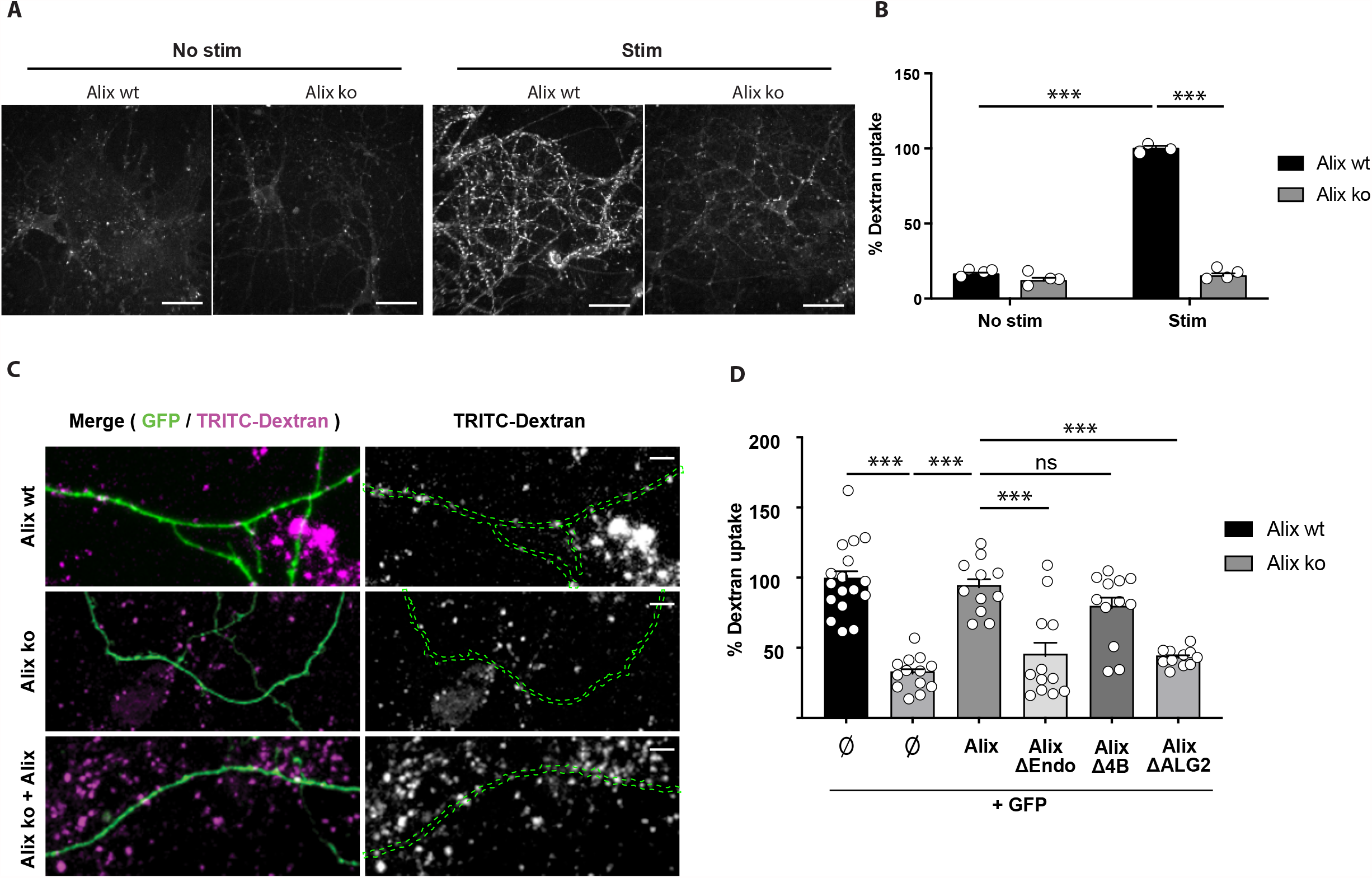
Alix-driven bulk endocytosis requires its binding to ALG-2 and endophilin but not to CHMP4. **A**) Confocal images of Alix wt and Alix ko hippocampal neurons stimulated with Bicucculine/4AP in the presence of 10kDa dextran. Scale bar: 50 μm. **B**) Dextran uptake triggered by stimulation is strongly reduced in Alix ko neurons. **C**) Representative images of Dextran uptake by GFP-expressing Alix wt and Alix ko neurons (Alix wt, top and Alix ko,middle) or Alix ko neurons expressing both GFP and Alix (Alix ko+Alix, bottom). Scale bars: 10 μm. **D**) Dextran uptake is rescued in ko neurons expressing Alix wt and AlixΔChmp4B (AlixΔ4B), but not AlixΔALG-2 or AlixΔendo. % of dextran uptake corresponds to the number of dextran spots per μm expressed as percentages of the control (Alix wt) for each experiment. <u>Average +/-SEM, N, statistical analysis:</u> **B)** 16.26+/-1.08; 11.86+/-2.05; 100+/-1.76; 15.10+/-1.76 for Alix wt no stim, Alix ko no stim, Alix wt stim, Alix ko stim respectively. N=4 experiments, p=0.0001, one-way ANOVA. **D)** 100+/-6.31; 33.27+/-3.32; 94.78+/-5.81; 45.89+/-9.36; 80.05+/-7.32; 44.59+/-1.85 for Alix wt, Alix ko, Alix ko + Alix, Alix ko + AlixΔChmp4B, Alix ko + AlixΔendo, Alix ko + AlixΔALG-2. N=17 neurons for Alix wt, n=13 neurons for Alix ko, n=11 for Alix ko + Alix, n=12 for Alix ko + AlixΔChmp4B, n=12 neurons for Alix ko + AlixΔendo,, and n=11 for Alix ko + AlixΔALG-2, all from 3-4 independent experiments, p=0.62 (Alix wt vs Alix AlixΔ4B) p=0.0001 for the other conditions, one-way ANOVA).

**Figure 5.**
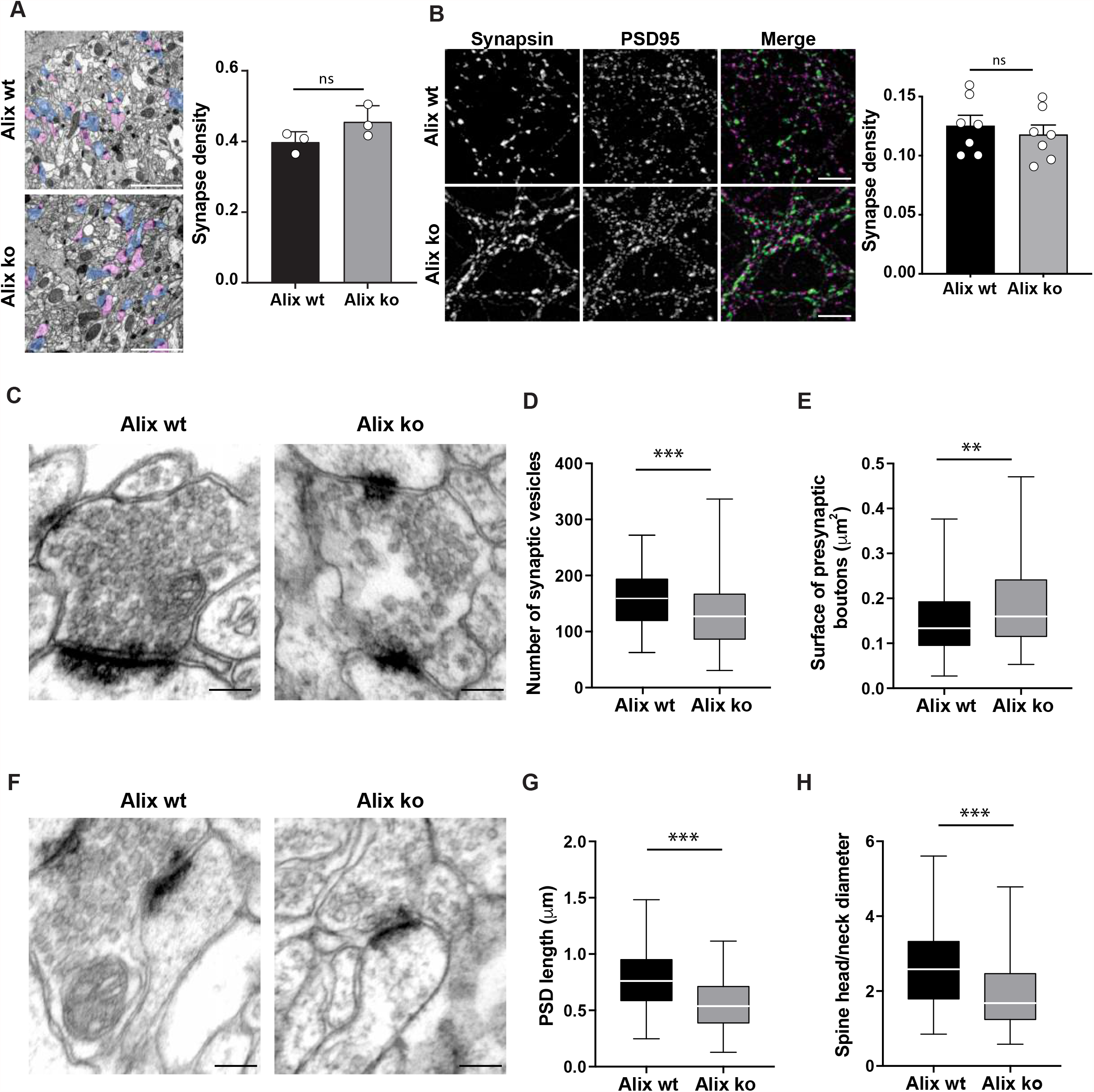
Morphological anomalies of Alix ko synapses. **A**) Representative electron micrographs of the CA1 stratum radiatum from Alix wt and ko brain sections. Presynaptic profiles are highlighted in blue, and dendritic spines are in purple. Scale bar: 2 μm. Graph shows no difference in synaptic density per μm^2^ of Alix wt and ko brains. **B**) 15 DIV hippocampal neurons were stained with anti-PSD95 (magenta) and anti-synapsin-1 (green) antibodies. Immunolabelled objects were considered as synapses when both stainings were juxtaposed. Scale bar: 10 μm. Graph shows no difference in the number of synapses per μm^2^ of Alix wt and ko neurons). **C-H**) Anomalies can be detected in synapses of Alix ko brains. **C, F**) Representative electron micrographs of CA1 from Alix wt and ko mice. Scale bar: 200 nm. Graphs represent the numbers of synaptic vesicles per μm^2^ (**D**), presynaptic bouton surface area (**E**), PSD length (**F**), ratio between the diameter of the spine head and neck (**G**). <u>Average +/-SEM, N, statistical analysis:</u> **A)** 0.40+/-0.02; 0.45+/-0.03 for Alix wt and Alix ko respectively. N=600 synapses from 3 animals per genotype, p=0.1395, Unpaired t-test. **B)** 0.13+/-0.01; 0.12+/-0.01 for Alix wt and ko respectively. N=7 independent experiments, p=0.5501, Unpaired t-test. <u>Median (min to max), N, statistical analysis</u> **D**) 165.4 (69 to 278); 127.3 (31 to 336) for Alix wt and Alix ko respectively. N=136 synapses from 3 animals, p=0.0009, Mann Whitney test). **E)** 0.13 (0.027 to 0.376); 0.16 (0.053 to 0.47) for Alix wt and Alix ko respectively. N=136 synapses from 3 animals, p=0.0036, Mann Whitney test). **G)**: 0.76 (0.25 to 1.48) ; 0.54 (0.13 to 1.11) for Alix wt and Alix ko respectively. N=136 synapses from 3 animals, p=0.0001, Mann Whitney test). **H)** 2.58 (0.85 to 5.60); 1.68 (0.58 to 4.78) for Alix wt and Alix ko respectively. N=136 synapses from 3 animals, p=0.0001, Mann Whitney test.

**Figure 6.**
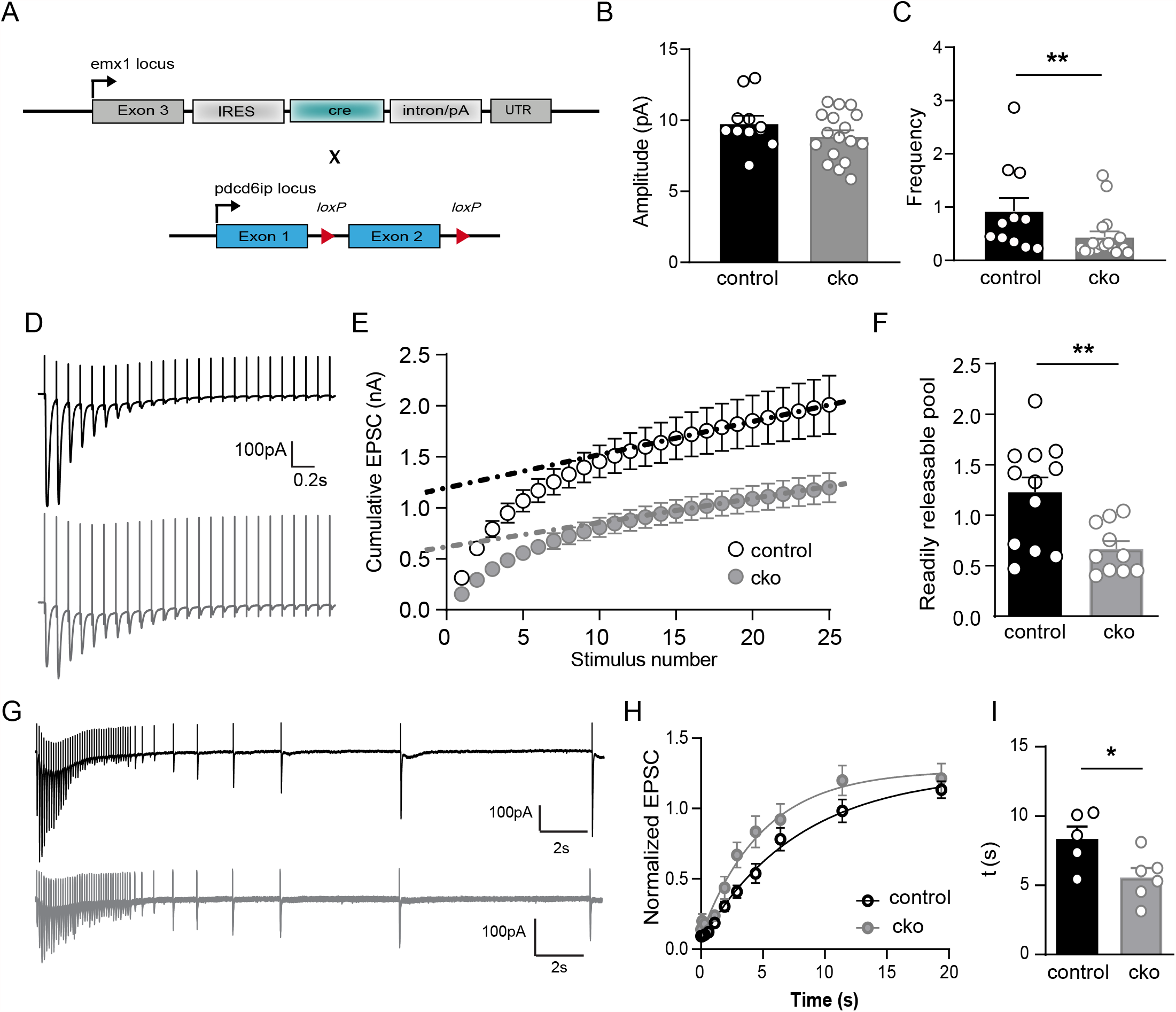
Voltage-clamp recordings uncover anomalies at Alix ko synapses. **A**) Emx1 ^IREScre^ (Emx-Cre) and Alix ^fl/fl^ mouse lines were crossbred to delete Alix in neocortical and hippocampal excitatory neurons (Alix cko). **B**) No difference in spontaneous Excitatory Post Synaptic Current (sEPSC) amplitude was detected between control and cko mice. **C**) The frequency of sEPSC in cko neurons is lower compared to control. **D**) Representative traces showing short-term depression in response to 10 Hz stimulation trains in control (black) and cko (grey) mice. **E**) Cumulative EPSC amplitudes in response to 10 Hz stimulation. Train-extrapolation is illustrated by the dashed line. **F**) The size of the readily releasable pool, estimated by the train-extrapolation method, was significantly decreased in Alix cko mice. **G**) Representative traces showing recovery from depression in response to 10 Hz stimulation trains in control (black) and Alix cko (grey) mice. **H**) To evaluate the speed of recovery, all inward signals were normalized to 1^st^ inward current. Recovering signals with a single-exponential function revealed the plateaus at which the capacities of recovery were saturated. **I**) Alix cko neurons slightly recover faster than controls. <u>Average +/-SEM, N, statistical analysis:</u> **B)** 9.79+/-0.53; 8.88+/-0.42 for controls and Alix cko respectively. N=11 neurons from 5 control mice and N=17 neurons from 6 Alix cko mice, p=0.1875, unpaired t test. **C)** 0.93+/-0.25; 0.44+/-0.10 for controls and Alix cko respectively. N=11 neurons from 5 control mice and N=17 neurons from 6 Alix cko mice, **p=0.0095, Mann Whitney test. **F)** 1.23+/-0.15; 0.67+/-0:08 for controls and Alix cko respectively. N=12 neurons from 5 control mice, and N=10 neurons from 4 Alix cko mice, **p=0.0052, unpaired t test. **I)** 8.34+/-0.90; 5.56+/-0.69 for controls and Alix cko respectively. N= 5 and 6 neurons from 3 mice from control and Alix cko respectively, *p=0.0336, unpaired t test.

**Figure 7.**
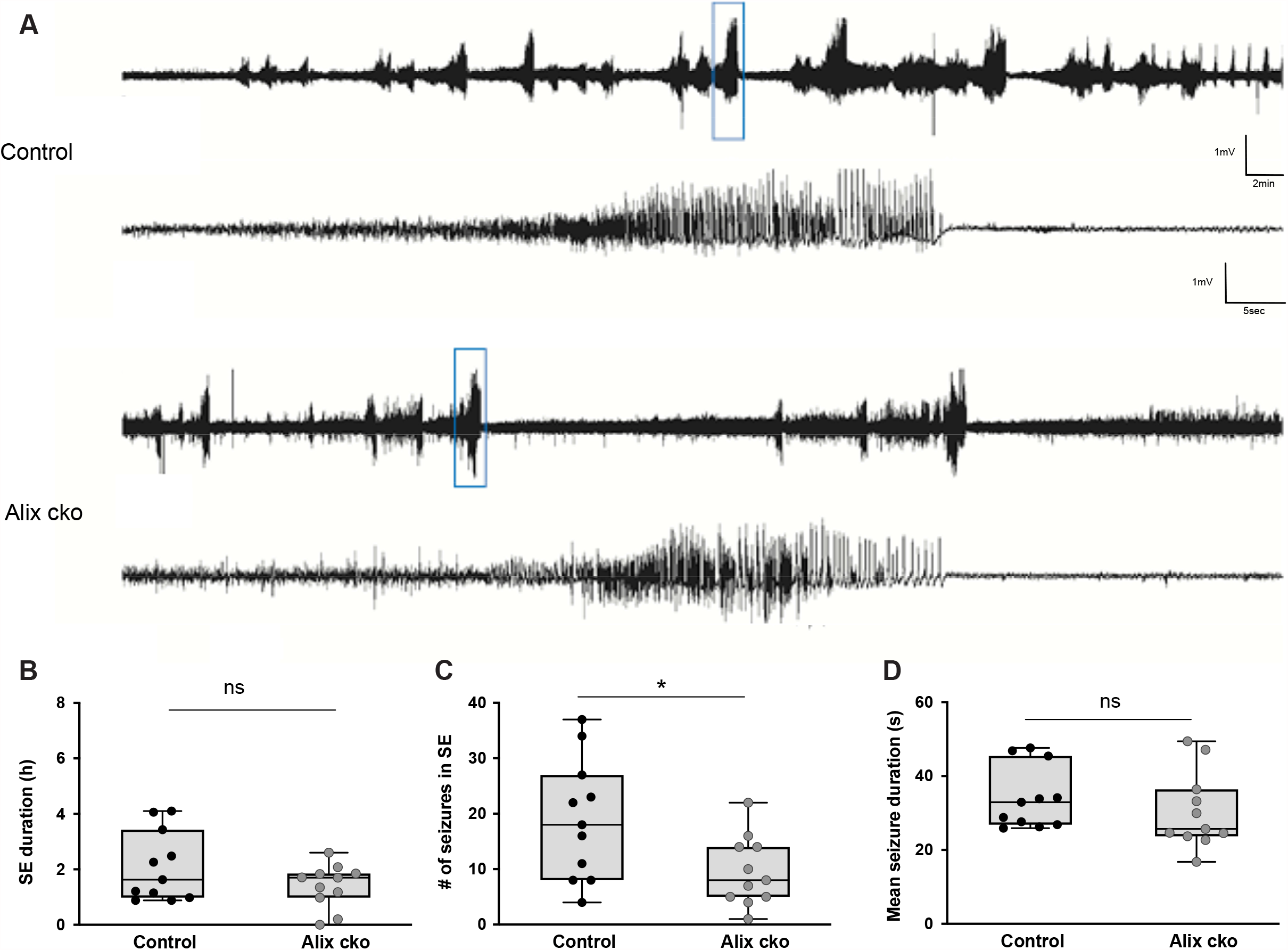
Alix cko mice develop less seizures during status epilepticus. **A**) Representative EEG traces from control and Alix cko mice. **B**) Total duration of *status epilepticus* (SE) did not differ between Alix cko and control mice. **C**) Alix cko mice experience about 66 % less seizures during SE than controls. **D**) Mean seizure duration during SE was not affected in Alix cko mice compared to controls. <u>Median + IQR, N, statistical analysis:</u> **A, B**) 1.6 h (IQR = 2.5); 1.7 h (IQR = 0.9) for control and Alix cko mice respectively. N=11 mice for both genotype, p= 0.49, Mann-Whitney test. **C**) 18 seizures (IQR = 9); 8 seizures (IQR = 9) for control and Alix cko mice respectively. N=11 mice for both genotypes, *p= 0.03, Mann-Whitney test. **D**) 32.9 s (IQR = 18.6); 25.7 h (IQR = 12.7) for control and Alix cko mice respectively. N=11 mice for both genotype, p= 0.19, Mann-Whitney test.

### Synaptic anomalies detected in Alix ko mice

To study the effect of Alix deletion *in vivo*, we first compared brains of Alix ko (Laporte et al., 2017) and wt mice and found that the density of synaptic contact as well as dendritic spines in the CA1 hippocampal region of adult brains was not different (**Figure 5A, Supplementary Figure 4**). This indicates that Alix is not required for synaptogenesis, a conclusion confirmed *in vitro* since the number of synapses revealed by co-immunostaining with synapsin-1 and PSD95 was similar between wt and Alix ko cultured hippocampal neurons (**Figure 5B**). However, examination of the CA1 *stratum radiatum* of adult mice showed that Alix ko synapses contained significantly fewer synaptic vesicles (SV) than wt (**Figure 5C, D**). Furthermore, the size of postsynaptic densities, known to be strictly correlated with the number of SV (Harris and Stevens, 1989), was similarly reduced in Alix ko synapses (**Figure 5F, G**). Importantly, the surface of Alix ko synaptic boutons was also significantly increased possibly suggesting plasma membrane accumulation in these synapses (**Figure 5C, E**). Finally, at the postsynaptic level, we also noticed that the ratio between the diameter of the spine head and that of the neck was changed in Alix ko animals, which can reflect defects in maturation or plasticity of adult synapses (**Figure 5F, H**). Accordingly, field recordings in the CA1 *stratum radiatum* of hippocampal slices revealed that long-term potentiation (LTP) induced by tetanic stimulations was significantly impaired in Alix ko hippocampal slices (**Supplementary Figure 5**).

### Voltage-clamp recordings uncovers anomalies at Alix ko synapses

In order to better characterize the effect of the lack of Alix on the physiology of synapses, we next used Alix conditional-ko mice where Alix is deleted in neocortical and hippocampal excitatory neurons (Gorski et al., 2002) (**Figure 6A**). We first recorded CA1 neurons of hippocampal slices and found that the amplitude of spontaneous EPSCs was unaffected whereas the frequency of events was significantly lower in Alix ko neurons (**Figure 6B, C**). This was reminiscent of the reduced frequency of burst measured by MAE in dissociated hippocampal cultures (**Supplementary figure 3C, D**). Stimulation of Schaffer collaterals revealed that evoked EPSCs have a smaller amplitude in Alix ko brains than in wt animals (**Figure 6D, E**). In order to estimate the amount of the readily releasable pool (RRP), we performed repeated stimulations and integrated the EPSC amplitudes to give a cumulative plot during 10 Hz trains. Linear regression fit to the last 10 data points was back extrapolated to time 0 (dotted line) to estimate the cumulative EPSC before steady state depression. This rough estimate of the RRP size (Schneggenburger et al., 1999) revealed that it is two times smaller in Alix ko neurons (**Figure 6F**), in good agreement with the lower amounts of SVs quantified in EM sections of synapses of the CA1 *stratum radiatum* (**Figure 5D**). Interestingly, we could observe that the rate of recovery post-depression appears faster in Alix ko mice, a possible mechanism of compensation for the reduced pool of synaptic vesicles (**Figure 6G, H, I**).

Thus, these electrophysiological results show that synapses of Alix ko neurons which lack ABDE have less SVs but increased rate of endocytosis, as already suggested in cultured neurons by the increase of Syp-pH endocytosis and of newly formed SVs detected in Alix ko-hippocampal and CB neurons, respectively.

### Alix conditional ko mice undergo less kainate induced acute seizures

We next examined the possible *in vivo* consequences of synaptic changes detected during repeated stimulation of Alix ko neurons in hippocampal slices. Because of our earlier finding of Alix increase at synapses of the rat hippocampus after kainaite injections (Hemming et al., 2004) we tested how the lack of Alix might influence kainate induced epileptic seizures, a mouse model for human temporal lobe epilepsy (Bedner et al., 2015). The unilateral intracortical kainate injection induces the acute phase, *status epilepticus*, characterized by high seizure frequency, during several hours. Seizure activity was determined by telemetric EEG recording and synchronized video monitoring (**Figure 7A**). Here we used again the conditional-ko mice where Alix is deleted in neocortical and hippocampal excitatory neurons (**Figure 6A**). The total duration of *status epilepticus* (SE) was comparable in Alix ko and control mice (**Figure 7B**). However, Alix cko mice experienced about 70 % less seizures than control mice (**Figure 7C**). The average seizure duration was only minimally reduced in Alix ko compared to control (**Figure 7D**). These results show that the absence of Alix selectively reduces the number of high frequency events (single seizures), without affecting the total duration of *status epilepticus*.

This kainate injection paradigm produces seizures originating from the hippocampus, which can spread to other contralateral brain areas (Bedner et al., 2015). Microglial reactivity was detected 24 h after kainate injection in ipsilateral and contralateral cortex and hippocampus in both control and Alix cko animals (**Supplementary Figure 6A-E**). While the difference between ipsi-and contralateral side was non-significant in control animals (≈ 10%), Alix ko animals displayed a significantly reduced contralateral microglial activation (≈ 30%) (**Supplementary Figure 6E**). No difference between Alix ko and control animals was detected using GFAP immunolabelling, an expected result since astroglial activation peaks 4-7 days after kainate injection (**Supplementary Figure 6F**). Thus, the reduced contralateral microglial activation seen in Alix ko animals indicates a reduced propagation of the seizure activity and corroborates the previous seizure quantification (**Figure 7C**).

In summary, Alix is specifically required for ADBE. Impairment in this process seems to relate to impairments in synaptic function and plasticity in normal and pathological settings.

## Discussion

To sustain neurotransmission and prevent expansion of the presynaptic plasma membrane, synaptic vesicle (SV) fusion is coupled to the endocytic recycling and regeneration of SV proteins and lipids. Vesicle components can be retrieved from the plasma membrane via clathrin scaffolds, or via clathrin-independent processes mediating fast and ultrafast endocytosis and, in the case of high frequency stimulation, bulk endocytosis (Gan and Watanabe, 2018). Using Alix ko cells, we recently discovered that Alix drives clathrin-independent-endocytosis (CIE) during ligand-induced endophilin-dependent endocytosis as well as bulk endocytosis (Laporte et al., 2017; Mercier et al., 2016). We had previously reported that Alix immunoreactivity in the rat hippocampus is strongly upregulated in synapses undergoing high frequency activation during kainate-induced epileptic seizures. This increase was presynaptic and only transient, as it was reversed soon after cessation of the seizures (Hemming et al., 2004). The aim of the present study was to decipher the role of Alix in endocytosis at synapses. We first used primary cultures of cortical and hippocampal neurons, which make networks undergoing spontaneous activities. Massive increases in spike frequency induced by 4AP and bicuculline resulted in elevated calcium entry (Hardingham et al., 2001) and triggered activity dependent bulk endocytosis (ADBE) together with a transient concentration of Alix at presynaptic parts. Quantifying bulk endocytosis using dextran uptake or electron microscopy observations showed that ADBE is strongly reduced in Alix ko neurons demonstrating that Alix is essential for ADBE. ADBE is triggered by high [Ca^2+^] in response to sustained activity (Paillart et al., 2003). Calcium and calmodulin-dependent phosphatase calcineurin were suggested to make a link between synaptic activity and formation of bulk endosomes through dynamin (Morton et al., 2015). Alix may represent another link as its recruitment and activity in ADBE seems dependent on calcium increase. Indeed, EGTA, which binds calcium at an approximately 100 times slower on-rate than BAPTA (Adler et al., 1991; Schneggenburger and Neher, 2005), and was already known to block ADBE without affecting synaptic release, was equally efficient as BAPTA in inhibiting Alix recruitment. Furthermore, Alix recruitment required its capacity to bind the calcium-binding protein ALG-2, which also concentrated at hyperactive synapses. ALG-2 is a cytosolic penta-EF hand-containing protein with two Ca^2+^ binding sites (Kd= 1.2 µM), and its interaction with Alix strictly depends on calcium (Missotten et al., 1999; Shibata et al., 2004; Trioulier et al., 2004). Conformational change and exposure of hydrophobic patches occur at µM concentrations of Ca^2+^, suggesting that ALG-2 functions as a calcium sensor (Maki et al., 2016). Indeed, in non-neuronal cells, calcium entry provoked by membrane wounds leads to the sequential recruitment to the membrane of ALG-2, Alix and ESCRT-III proteins necessary for membrane repair (Jimenez et al., 2014; Scheffer et al., 2014). In neurons, the activity-dependent accumulation of ALG-2 at synapses required Ca^2+^ binding as it was totally abolished by point-mutations within the two Ca^2+^ binding sites. On the other hand, it did not require Alix, as ALG-2 recruitment also occurred in Alix ko neuron. Thus, our observations highlight synaptic ALG-2 as an obvious candidate for sensing calcium elevation and suggest the following scenario: sustained high frequency depolarization leads to massive elevation of calcium in the synaptic bouton; Ca^2+^ binding to ALG-2 causes its accumulation at the synaptic membrane and binding to cytosolic Alix, whose recruitment to the membrane drives ADBE.

Little is known about the molecular mechanisms underlying the plasma membrane deformation during ADBE. The proline rich domain of Alix interacts with the SH3 domains of endophilins-A1-3 which contain N-BAR domains and have been shown to regulate clathrin-dependent and -independent SV endocytosis at different synapses (Gad et al., 2000; Llobet et al., 2011; Ringstad et al., 1999). BAR domains are dimerization domains able to induce, stabilize and sense membrane curvature (Farsad et al., 2001; Kjaerulff et al., 2011). Interestingly, a proteomic approach revealed the presence of endophilin-A1, Alix and ALG-2 in bulk endosomes (Kokotos et al., 2018). Furthermore, endophilin-A2, actin and dynamin mediate a restricted type of CIE activated upon ligand binding to cargo receptors (Boucrot et al., 2014; Renard et al., 2014). We previously showed that in fibroblasts, endophilin-A and Alix act in the same CIE pathway, even if Alix is more promiscuous for ligands, as it is also involved in fluid phase endocytosis. In this case interaction of Alix with endophilins favoured their presence at the membrane (Mercier et al., 2016). We now found that endophilin-A2 decorated presynapses of resting neurons and that Alix was needed for its synaptic enrichment during stimulation. An Alix point mutant unable to interact with endophilin was recruited to synapses but could not rescue ABDE in Alix ko neurons, reminiscent of the situation seen in fibroblasts (Mercier et al., 2016). Thus, a complex between endophilin-A and Alix is required for ADBE. One function of Alix might be to enable endophilin-A binding onto flat membranes of the peri-active zone to drive membrane bending in response to high calcium concentrations (Boucrot et al., 2014; Kjaerulff et al., 2011). Interestingly, the incapacity of Alix to interact with Chmp4B from ESCRT-III, known to alleviate its plasma membrane repair ability (Jimenez et al., 2014; Scheffer et al., 2014), did not alter rescue of ADBE. This result also discriminates Alix-driven ADBE from roles of the ESCRT system in the degradation of SV proteins (Sadoul et al., 2017; Sheehan et al., 2016).

Electrophysiological recordings and EM observations in CA3-CA1 synapses both point to a reduction in the number of synaptic vesicles due to the lack of Alix and possibly to the associated defects in ADBE. On the other hand, Syp-pH fluorescence and EM analysis revealed that activity dependent synaptic endocytosis is increased in absence of Alix. Accordingly, electrophysiological patch-clamp recordings show that Alix ko CA3-CA1 synapses recover better from STD induced by repeated stimulations Thus, compensation by increased endocytosis, is not sufficient to rescue the phenotype. This could be a first demonstration that ADBE of the essential role of ADBE in regulating the number of synaptic vesicles and thereby normal synaptic function.

Excitatory, glutamatergic neurotransmission plays a central role in the generation of seizure activity (Albrecht and Zielińska, 2017; Barker-Haliski and White, 2015) and figures among the primary anti-epileptic drug targets (Rogawski and Löscher, 2004). Sustained, intense and synchronous neurotransmission as occurring during seizures requires replenishment of synaptic vesicle pools mainly through activity-dependent bulk endocytosis (ADBE) (Cheung and Cousin, 2012). Impairing ADBE *in vivo*, by deleting Alix in neocortical and hippocampal excitatory neurons led to a significantly reduced number of seizures during SE, without affecting the total duration of SE or the mean seizure duration. The present results suggest that excitatory networks lacking Alix have a reduced capacity to initiate sequential seizures, possibly due to an impaired replenishment of synaptic vesicles through ADBE. Seizure termination and *status epilepticus* cessation however seem to be governed by distinct mechanisms, probably involving inhibitory transmission to re-establish the balance between excitation and inhibition in the brain. Our results highlight Alix ko mice as an invaluable tool for exploring and understanding the exact role of ADBE at synapses undergoing normal and pathological stimulations.

## Acknowledgements

We thank J. Brocard for helping with data analysis, Y. Saoudi for helping with fluorescence microscopy, N. Liaudet from the Bioimaging core facility of the University of Geneva for his help with the segmentation and electron microscopy, F. Saudou for making the MEA experiments possible. We also thank K. Sadoul for critically reading the manuscript. This work was funded by France Alzheimer (R.S.), Ministère de l’Enseignement Supérieur et de la Recherche (K.I.C, M.R. and M.H.L.), Marie Sklodowska-Curie post-doctoral fellowship (M.M.); EC H2020 MSCA-ITN EU-GliapHD #722053 (L.C.C. and F.K.), Agence Nationale de la Recherche ANR (J.M.-H.) and Fondation pour la Recherche Medical FRM (E.M.).

## Author contributions

RS conceived the project. MHL participated to most of the in vitro experiments and analysed electron microscopy data under the supervision of SF. KIC performed and analysed most of the *in vitro* experiments concerning live imaging of Alix recruitment and dextran uptake. LCC performed and analysed the EEG telemetry and unilateral intracortical kainate injection. NZ performed and analysed patch clamp electrophysiological recordings. The experiments of LCC and NZ were done under the supervision of FK. MR and FLa performed field electrophysiological recordings. JM-H performed electron microscopy of mouse brains. MM and DP performed and analysed the live imaging of synaptophysin-pHluorin during electrical stimulation. CC performed mouse breeding, genotyping and participated in *in vitro* experiments. ED helped with data analysis of live imaging. VM performed FIB-SEM imaging. BB performed mouse genotyping and construction of plasmids. EM and MC performed and analysed multielectrode array experiments. FLe did some of the live imaging experiments of Alix recruitment *in vitro*. FJH performed electron microscopy of cultured neurons. RS and MHL wrote the manuscript with input from all other authors.

