## Supplementary figures and legends for "ALG-2 interacting protein-X (Alix) is required for activity-dependent bulk endocytosis at brain synapses"

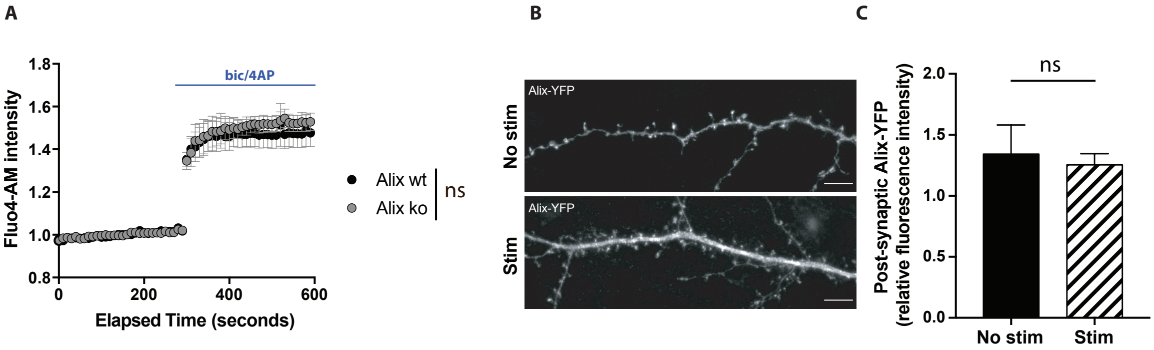


**Supplementary figure 1. A**) No difference in calcium rise in Alix wt and ko hippocampal neurons upon bicuculline/4AP stimulation. Fluo4-AM intensity corresponds to the change in fluorescence, normalized to initial fluorescence (N=4 experiments, p=0.9999, two-way ANOVA). **B**, **C**) Alix-YFP does not accumulate in dendritic spines upon bicuculline/4AP stimulation of 15DIV hippocampal neurons. Scale bar: 5 µm. Postsynaptic Alix-YFP corresponds to the ratio between YFP-fluorescence at PSD95 labelled ROI and at neighbouring dendritic parts.

<u>Average +/- SEM, N, statistical analysis</u>: 1.34+/- 0.24; 1.25+/- 0.09 for no stim and stim respectively. N=33 and n=25 neurons for both conditions, from 5 experiments, p=0.3821, Mann Whitney test).

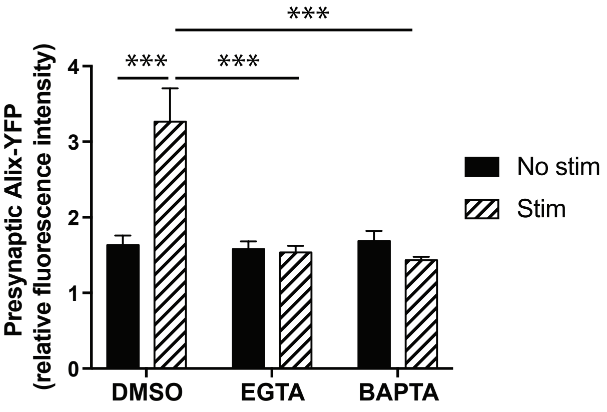


**Supplementary figure 2.** Calcium chelators (BAPTA-AM and EGTA-AM) block presynaptic Alix recruitment upon bicuculline/4AP stimulation. Presynaptic Alix-YFP corresponds to the ratio of fluorescence between presynaptic and non-synaptic axonal ROI.

<u>Average +/- SEM, N, statistical analysis</u>: 1.63+/- 0.12; 3.27+/- 0.44; 1.58+/- 0.10; 1.54+/- 0.009; 1.69+/- 0.13; 1.43+/- 0.04 for DMSO no stim, DMSO stim, EGTA no stim, EGTA stim, BAPTA no stim, BAPTA stim respectively .N=12, 9 and 9 neurons for DMSO, EGTA and BAPTA respectively. p=0.0001, one-way ANOVA).

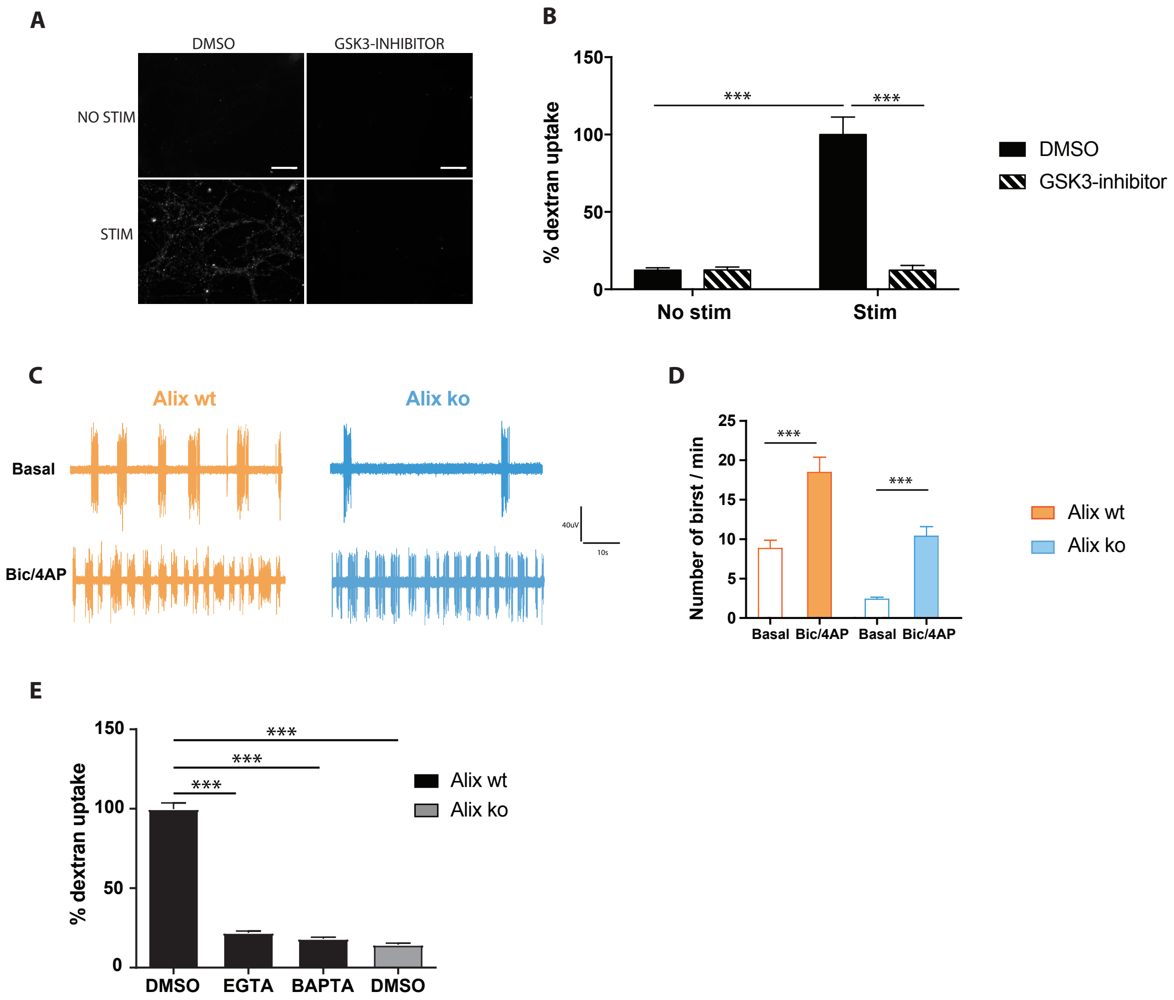


**Supplementary figure 3. A**, **B**) Dextran uptake is abolished in Alix wt neurons treated with an inhibitor of bulk endocytosis (GSK3-inhibitor). Confocal images of Alix wt hippocampal neurons stimulated in the presence of 10 kDa dextran with or without a GSK3 inhibitor. Scale bar: 50 µm. % dextran uptake corresponds to the number of dextran spots per ROI expressed as percentages of the positive control **C**, **D**) Multiple Electrode Array activity recordings of 15 DIV hippocampal neuron cultures showing the effect of bicucculine/4AP incubation for 10 min. Representative traces of wt (orange) and ko (blue) cultures are shown on panel **C**.**E**) Dextran uptake is abolished in wt neurons by calcium chelators EGTA and BAPTA. % dextran uptake corresponds to the number of dextran spots per ROI expressed as percentages of the positive control for each experiment.

<u>Average +/- SEM, N, statistical analysis:</u> **B**) 12.41+/- 1.51; 100+/- 11.29; 12.46+/- 1.90; 12.40+/- 2.99 for DMSO no stim, DMSO stim, GSK3-inhibitor no stim, GSK3-inhibitor stim respectively. N=3 experiments, p=0.0001, one-way ANOVA. **D**) 12.41+/- 1.51; 100+/- 11.29; 12.46+/- 1.90; 12.40+/- 2.99 for DMSO no stim, DMSO stim, GSK3-inhibitor no stim, GSK3-inhibitor stim respectively. N=3 experiments, p=0.0001, one-way ANOVA). D) average +/- SEM are as follow: 8.81+/-1.05; 18.48+/-1.92; 2.41+/-0.27; 10.33+/-1.23 for Alix wt no stim, Alix wt stim, Alix ko no stim, Alix ko stim respectively. N=53 and 58 field of view for Alix wt and Alix ko respectively, p<0.0001, one-way ANOVA. **F**) 100+/- 0; 21.89+/- 4.19; 18.34+/- 3.54; 14.53+/- 4.17.N=3 experiments, p=0.0001, one-way ANOVA.

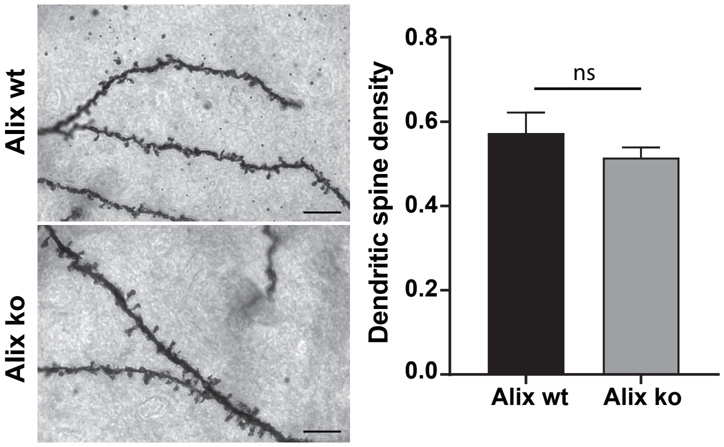


**Supplementary figure 4.** Brain sections from 8-week-old Alix wt or ko were stained by the Golgi-Cox impregnation technique. Stained dendritic segments were visualized by bright-field microscopy. Scale bar: 10 μm. Numbers of spines per μm of dendrites were counted.
<u>Average +/- SEM, N, statistical analysis:</u> 0.57 +/- 0.23; 0.51+/-0.01 for Alix wt and Alix ko respectively. N=3 animals per genotype, p=0.1414, Unpaired t test).

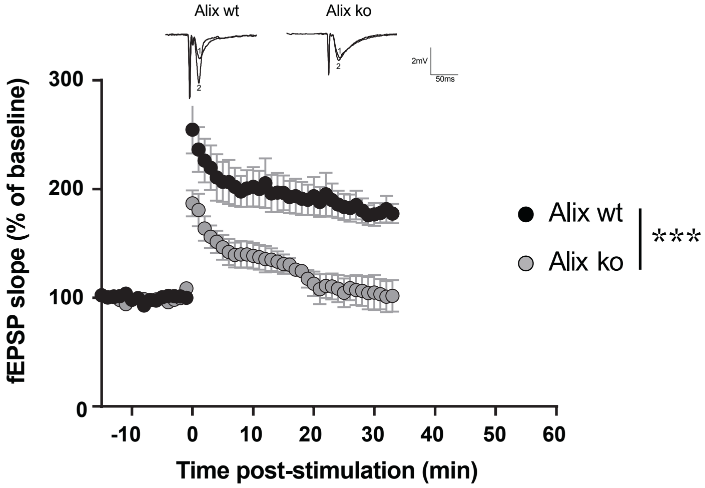


**Supplementary figure 5.** Long-term potentiation of fEPSP slope evoked by high frequency stimulation of Schaffer collaterals delivered at time 0. Inserts show representative EPSPs traces.
<u>N, statistical analysis:</u> n=5 slices from 3 animals per genotype, p=0.0001, two-way ANOVA.

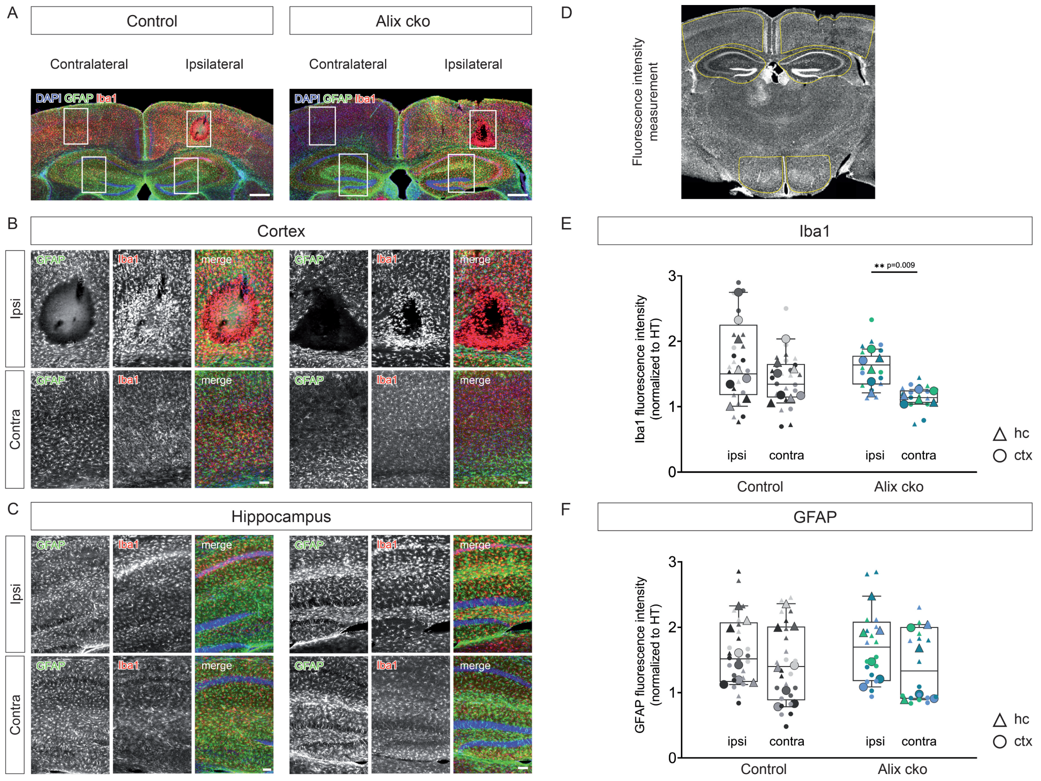


**Supplementary figure 6. A**) Coronal brain section from control or Alix cko stained for DAPI (blue), GFAP (green) and Iba1 (red). 24Scale bar= 500 µm. **B**, **C**) Magnification of highlighted cortical (**B**) and hippocampal (**C**) areas show an increased ipsilateral (ipsi) microglial activation (Iba1) compared to contralateral (contra), which was more pronounced in Alix ko mice. Astroglial reactivity (GFAP) was moderately increased adjacent to the injection site in both groups. Scale bars= 100 µm. **D**) Depiction of fluorescence intensity measurement areas in cortex, hippocampus and hypothalamus. **E**) In Alix ko mice, the contralateral Iba1 immunoreactivity was about 30 % reduced in comparison to ipsilateral, in contrast to control mice. **F**) GFAP immunoreactivities were not significantly different between hemispheres and experimental groups, as expected from the early analysis time point of only 24 h after kainate injection. Cortical GFAP immunoreactivity (**A**) was similar to hypothalamus (normalized fluorescence intensity ≈ 1). The hippocampal GFAP expression (**B**) showed a higher variability, however approximately twice as high (normalized fluorescence intensity≈ 2), compared to cortex and hypothalamus. These data reflect the regional heterogeneity in astroglial GFAP expression, acting as internal confirmation of the analysis method. Circles and triangles represent individual quantifications from cortex and hippocampus, respectively. Large data points correspond to the average of three slices from the same animal (small data points in the background, colour-coded per animal). Fluorescence intensity values were normalized to the hypothalamic area (HT) of the respective hemisphere.

Mean +/- IQR, N, statistical analysis: **E**) Control : 1.1 (0.2) vs. 1.6 (0.4), contra- vs. ipsilateral medians (IQR); **p= 0.009); Alix cko: 1.3 (0.5) vs. 1.5 (1.1), contra- vs. ipsilateral medians (IQR); p= 0645. N (control) = 4 animals, n (Alix ko) = 3 animals. **F**) control: 1.4 (1.1) vs. 1.5 (0.9), contra- vs. ipsilateral medians (IQR); p= 0.6; Alix ko: 1.3 (1.1) vs. 1.7 (0.9), contra- vs. ipsilateral medians (IQR); p= 0.6. N (control) = 4 animals, n (Alix ko) = 3 animals.
